# One-carbon unit supplementation fuels tumor-infiltrating T cells and augments checkpoint blockade

**DOI:** 10.1101/2023.11.01.565193

**Authors:** Xincheng Xu, Zihong Chen, Caroline R. Bartman, Xi Xing, Kellen Olszewski, Joshua D. Rabinowitz

**Affiliations:** Department of Chemistry, Princeton University.; Lewis-Sigler Institute for Integrative Genomics, Princeton University; Ludwig Institute for Cancer Research, Princeton Branch, Princeton University.

## Abstract

Nucleotides perform important metabolic functions, carrying energy and feeding nucleic acid synthesis. Here, we use isotope tracing-mass spectrometry to quantitate the contributions to purine nucleotides of salvage versus *de novo* synthesis. We further explore the impact of augmenting a key precursor for purine synthesis, one-carbon (1C) units. We show that tumors and tumor-infiltrating T cells (relative to splenic T cells) synthesize purines *de novo*. Purine synthesis requires two 1C units, which come from serine catabolism and circulating formate. Shortage of 1C units is a potential bottleneck for anti-tumor immunity. Elevating circulating formate drives its usage by tumor-infiltrating T cells. Orally administered methanol functions as a formate pro-drug, with deuteration enabling control of formate-production kinetics. In MC38 tumors, safe doses of methanol raise formate levels and augment anti-PD-1 checkpoint blockade, tripling durable regressions. Thus, 1C deficiency can gate antitumor immunity and this metabolic checkpoint can be overcome with pharmacological 1C supplementation.

**Statement of significance:** Checkpoint blockade has revolutionized cancer therapy. Durable tumor control, however, is achieved in only a minority of patients. We show that the efficacy of anti-PD-1 blockade can be enhanced by metabolic supplementation with one-carbon donors. Such donors support nucleotide synthesis in tumor-infiltrating T cells and merit future clinical evaluation.

## Introduction

Cancer metabolism was revitalized early this century by aspirations to cut off the tumor fuel supply. Intensive research efforts explored central carbon pathways feeding both energy and biosynthetic metabolism [1,2]. Identification of alternative nutrient acquisition routes based on nutrient scavenging [3,4], such as micropinocytosis [5,6], enriched the field but complicated efforts to starve tumor cells, and up to now there has been no clinical success in targeting central metabolism for cancer therapy.

With the rise of immunotherapy, an alternative opportunity emerged: to remove metabolic barriers to successful antitumor immune response [7]. A myriad of such potential barriers have been identified, including intratumoral competition for glucose [8–10], amino acids [11,12], and oxygen [13,14], and potential metabolic impairment of immune cells by cancer cell-derived products like lactate [15] or kynurenine [16], with inhibitors of kynurenine synthesis entering the clinic but showing limited benefits in human patients [17].

Historically, the most therapeutically impactful area of cancer and immune metabolism, dating back to Sydney Farber [18], regards nucleotide metabolism, including folate enzymes that generate 1C units required for purine and thymidine synthesis [19]. Inhibitors of nucleotide metabolism are both anticancer agents and immunosuppressants. Anti-purines such as azathioprine, 6-mercaptopurine, and 6-thioguanine are frequently used in the treatment of autoimmune disease [20]. Methotrexate, which targets the production by dihydrofolate reductase of the biologically active form of folate, tetrahydrofolate (THF), is a first-line agent for T cell leukemia [21] and rheumatoid arthritis [22]. The clinical importance of these agents as immunosuppressants argues for purine biosynthetic capacity being a key determinant of immune response.

*De novo* purine synthesis starts with glutamine-mediated amination of phosphoribosyl pyrophosphate (PRPP), followed by a series of steps incorporating glycine and two folate-bound 1C units, in the form of formyl-tetrahydrofolate (formyl-THF). The 1C unit in tetrahydrofolate can come from multiple precursors, including serine or free circulating formate. Activated T cells upregulate *de novo* serine synthesis and the mitochondrial enzymes catalyzing conversion of serine into formyl-THF: SHMT2 and MTHFD2 [23]. Impairment of MTHFD2 leads to failure of effector T cell function and a shift towards a suppressor state [24]. Mitochondrial formyl-THF is exported to the cytosol as free formate, which together with formate imported from circulation, gets converted by MTHFD1 to cytosolic formyl-THF fueling *de novo* purine synthesis. CD8^+^ T cells with the loss of *Mthfd1* are strongly depleted in tumors and spleen, supporting the important role of formate incorporation for anti-tumor immunity [25].

The product of *de novo* purine synthesis is IMP, which can be converted in two steps into either AMP or GMP. These purines can also be synthesized via salvage pathways, by reaction of hypoxanthine, adenine, or guanine with PRPP to yield IMP, AMP, or GMP. Further phosphorylation yields the key energy molecules and nucleic acid precursors ATP and GTP. While the nucleobases hypoxanthine, adenine, and guanine are the direct substrates for purine salvage, their corresponding nucleosides (inosine, adenosine, guanosine) may also be major players in the inter-organ exchange of purines [26–28]. Inborn errors in nucleoside metabolism manifest as severe combined immunodeficiency [29–31]. Inosine is a component of an immune booster used in some viral infections (inosine pranobex) [32,33] and has been suggested to be a microbiome-derived product that supports cancer immunotherapy efficacy [34].

Here we investigate the relative contributions of *de novo* and salvage purine synthesis to the key purine end product ATP, in normal tissues, bulk MC38 tumors, splenic CD8^+^ T cells, and tumor-infiltrating CD8^+^ T cells (TILs). We find that tumors and TILs synthesize a particularly high fraction of their ATP purine ring *de novo*. We further show that both serine and free circulating formate provide 1C units to support TIL purine synthesis, and that pharmacological boosting of circulating formate by administering methanol as a formate prodrug enhances anti-PD-1 immunotherapy.

## Results

### Abundances of circulating purines

We began by establishing isotope-tracing methods for the source of the purine ring in ATP and GTP *in vivo*. Serine is an established tracer for *de novo* purine synthesis, which feeds into the ring by generating 1C units and glycine, which together account for 4 of the 5 carbon atoms in purines (Fig. 1A). Infusion of [U-^13^C]serine accordingly probes *de novo* purine production [35,36].

**Fig. 1.**
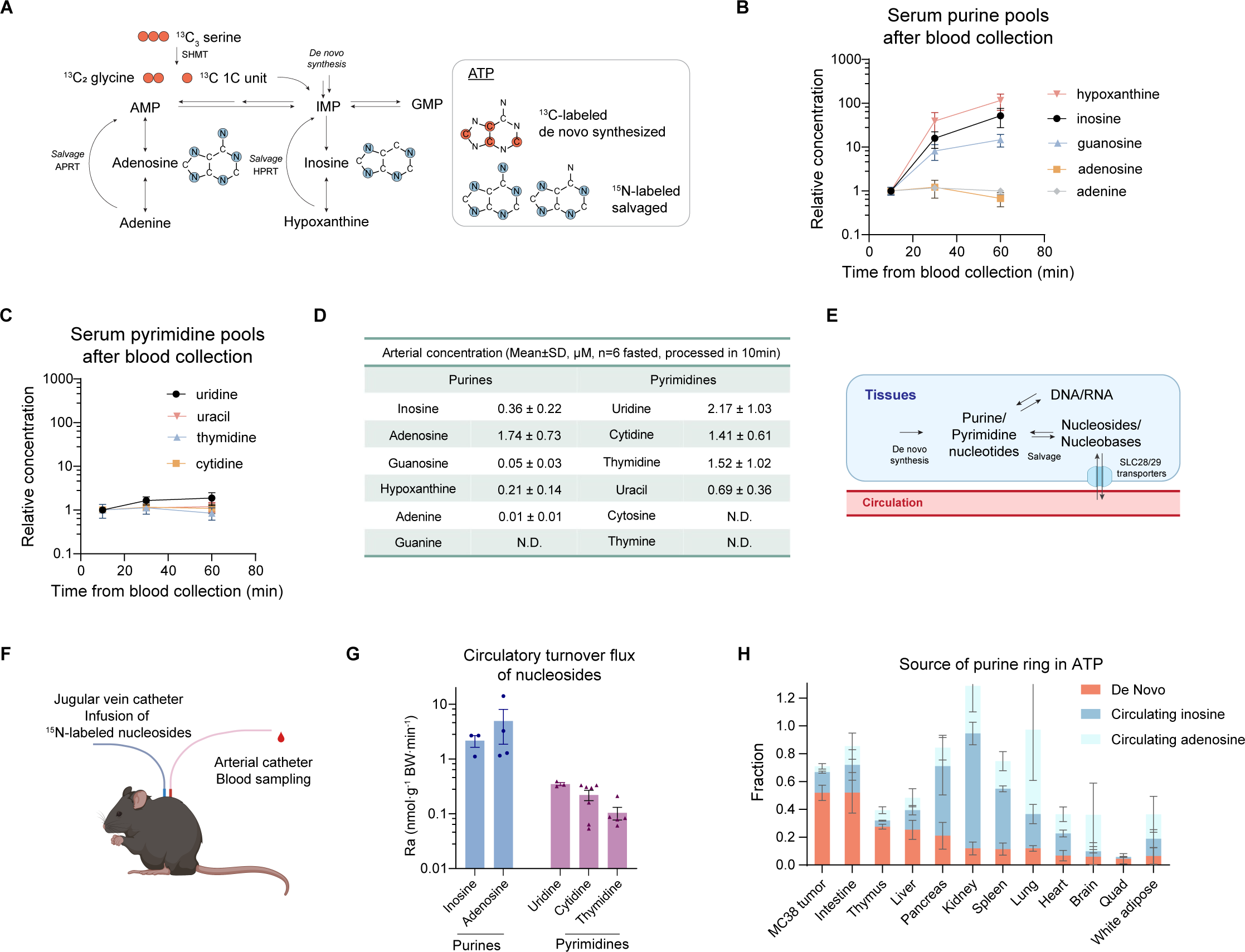
Source of ATP’s purine ring in tissues and tumors. (A) Schematic of tracing purine *de novo* synthesis and salvage using [U-^13^C]serine, [U-^15^N]inosine, and [U-^15^N]adenosine. (B) Serum concentrations of certain purine nucleosides and bases rise when isolated blood sits on ice (n = 3). (C) Serum concentrations of pyrimidine nucleosides and bases are relatively stable when isolated blood sits on ice (n = 3). (D) Concentrations of purine and pyrimidine species in quickly processed arterial serum from fasted mice (n = 6). Blood was kept on ice after sampling. Serum was isolated by centrifugation and extracted in 10 min from sample collection. N.D. = non-detectable. (E) Schematic diagram of nucleoside/nucleotide metabolism. (F) Schematic of measuring circulatory turnover flux (rate of appearance) of nucleosides in mice. (G) Circulatory turnover flux of purine and pyrimidine nucleosides. n = 3 for inosine; n = 4 for adenosine; n = 5 for uridine and thymidine; n = 6 for cytidine. (H) Contribution to ATP’s purine ring over 13 h tracer infusions by *de novo* synthesis (n = 3) and salvage of circulating inosine (n = 3) and adenosine (n = 3) in each tissue. Quad = quadriceps fermoris muscle. Unlabeled ATP reflects incomplete purine turnover during the infusion duration or production from other sources, likely primarily nucleic acid degradation. Mean ± SD except for mean ± SEM in (G); n indicates the number of mice.

Methods for measuring salvage synthesis were less well established. We began by carefully measuring the circulating concentrations of the potential salvage substrates. These measurements were technically complicated by the propensity for these low-energy metabolites to be rapidly produced by nucleotide degradation. For example, inosine and hypoxanthine concentrations in the tail blood were several-fold higher than in the arterial blood (which was sampled noninvasively from pre-catheterized mice) (Fig. S1A). Moreover, when blood was allowed to sit on ice, while most metabolite concentrations did not change, inosine and hypoxanthine increased greatly (Fig. 1B–C, Fig. S1B). Measurements from quickly-processed arterial serum showed undetectable purine nucleotides, and that inosine and adenosine are the most abundant circulating purines (and uridine, cytidine, and thymidine the most abundant circulating pyrimidines, Fig. 1D). Accordingly we focused on inosine and adenosine as tracers.

### Circulatory fluxes of nucleosides

We measured these nucleosides’ circulatory turnover flux (i.e. whole-body rate of release of these metabolites into the circulation, or “rate of appearance”, Fig. 1E) by infusing them in ^15^N-labeled form (Fig. 1F). This required continued care, as otherwise the labeling was diluted by unphysiological production during sampling (Fig. 1B, and Fig. S1B–D). We found that inosine and adenosine have much higher circulating fluxes than pyrimidine nucleosides (Fig. 1G).

[U-^15^N]inosine infusion labeled allantoin (a terminal purine degradation product, which plays a similar role in mouse to uric acid in human), to a similar extent as inosine itself (Fig. S1D). As allantoin labeling is insensitive to sampling method and sample handling (Fig. S1 D–E), we used its labeling as a surrogate for circulating inosine in subsequent experiments. Labeling of adenosine from inosine was limited (Fig. S1F), but not vice versa (Fig. S1G). This is consistent with inosine feeding into tissue nucleotides directly, whereas adenosine can contribute both directly and indirectly via circulating inosine, which can be produced from adenosine via the ubiquitous irreversible adenosine deaminase (ADA) reaction.

### Tumors synthesize purines *de novo*

We next explored the contributions of *de novo* synthesis ([U-^13^C]serine tracer) and salvage ([U- ^15^N]inosine and [U-^15^N]adenosine tracers) to tissue and MC38 murine colon cancer tumor purines by infusing these tracers for 13 h (Fig. 1A). The long infusion time is necessary for reaching isotopic pseudo-steady state due to the slow turnover of the purine skeleton relative to large tissue purine pools (Fig. S1H). While labeling differed depending on the tracer and target tissue, using either serine or inosine as the tracer, different purine nucleotides (IMP, AMP, ADP, ATP, GMP, and GDP) generally labeled to a similar extent, consistent with both serine and inosine entering all purines via the common precursor IMP (Fig. S1I–J). In contrast, in a subset of tissues, adenosine labeled adenosine nucleotides preferentially over guanosine nucleotides (Fig. S1 K). This reflects direct assimilation of adenosine without its prior conversion to IMP, as all five nitrogen atoms were retained (one is lost when adenosine becomes inosine) (Fig. S1L).

Because it is the most abundant purine, we focused on ATP as the most convenient readout. Inosine salvage, adenosine salvage, and *de novo* synthesis from serine each were the primary ATP purine ring contributors in certain tissues (Fig. 1H). For example, *de novo* synthesis predominated in intestine; salvage from inosine in pancreas, kidney, and spleen; and salvage from adenosine in lung. In MC38 tumors, *de novo* synthesis predominated.

### *De novo* purine synthesis in TILs versus splenic T cells

Tumors and organs are heterogeneous [37]. Given the importance of CD8^+^ T cells to anticancer immunity [38], we were particularly curious about the sources of purines in tumors versus splenic CD8^+^ T cells. In both tumors and immune organs (thymus, spleen), most salvage synthesis was from inosine (Fig. 1H). Accordingly, we focused on comparing tracing from serine versus inosine.

Using [U-^13^C]serine and [U-^15^N]inosine, ATP derived from serine versus inosine separate based on exact mass, allowing both tracers to be used in parallel in the same mouse. To isolate CD8^+^ T cells from mice receiving the dual tracers, we first dissociated normal spleen and MC38 tumors to single cells, followed by magnetic activated cell sorting (MACS) (Fig. 2A). The CD8^+^ enriched and depleted populations were sampled for flow cytometry validation (Fig. S2A–B), and the rest were placed directly into organic solvent to quench metabolism (Fig. 2A).

**Fig. 2.**
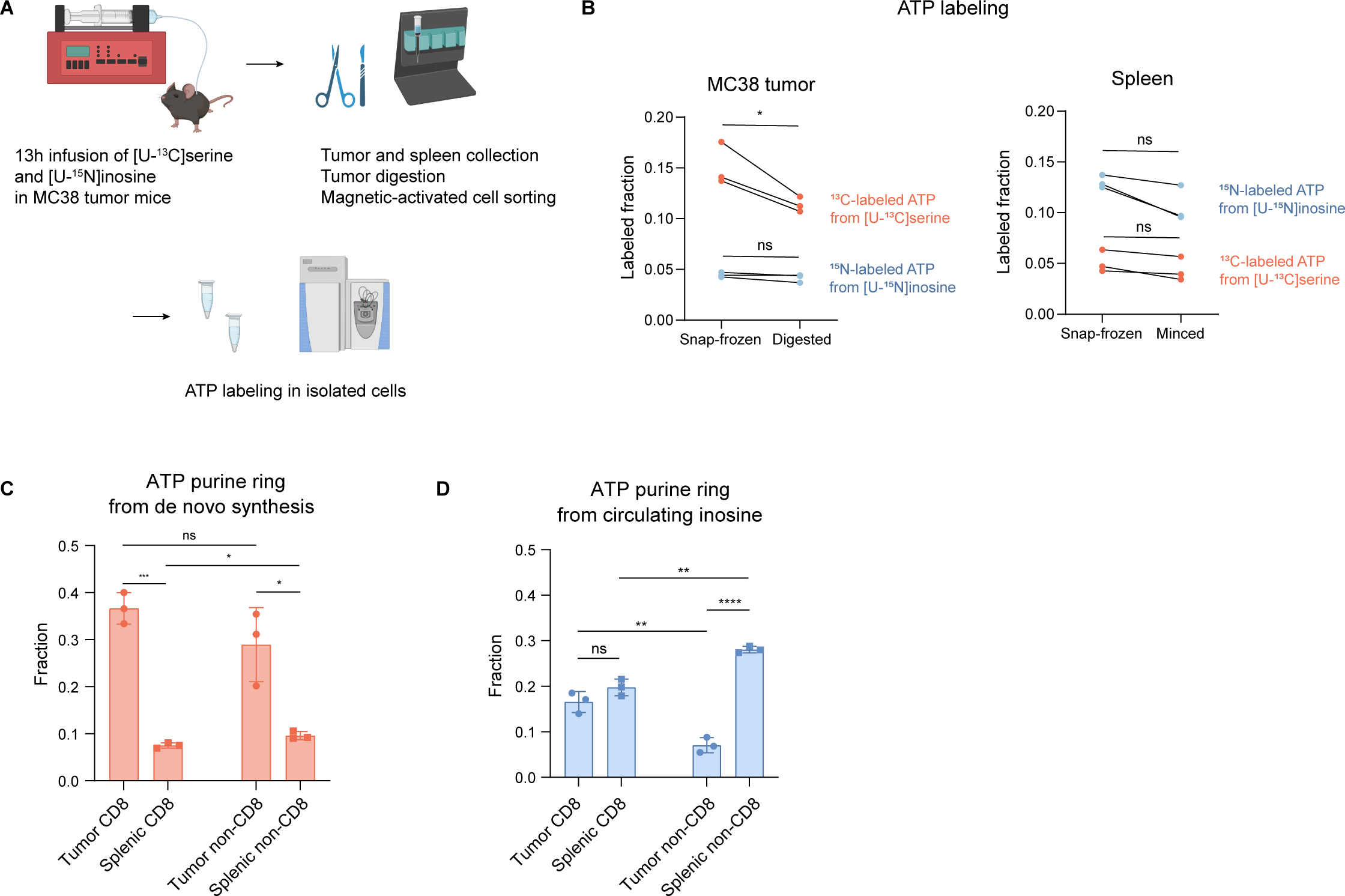
CD8^+^ TILs carry out more *de novo* purine synthesis than splenic CD8^+^ T cells (A) Experimental schema of tracing ATP source in different cell populations. Following 13-hour infusion of both [U-^13^C]serine and [U-^15^N]inosine (simultaneously) in mice bearing MC38 tumors, tumors were dissociated into single-cell suspensions. The CD8^+^ and CD8^-^ populations were fractionated by magnetic-activated cell sorting, and ATP labeling was measured by LC-MS. (B) The preservation of ATP labeling (from [U-^13^C]serine and [U-^15^N]inosine infusion) after tissue dissociation compared to the same sample being snap-frozen (n = 3). *P*-values were calculated by paired two-tailed Student’s *t*-test. (C–D) Fraction of ATP’s purine ring from *de novo* synthesis (C) versus circulating inosine in CD8^+^ or CD8^-^ populations in MC38 tumor and spleen. n = 3, mean ± SD. *P*-values were calculated by unpaired two-tailed Student’s *t*-test. n indicates the number of mice. ns, p > 0.05; * p < 0.05; ** p < 0.01; *** p < 0.001; **** p < 0.001.

A key challenge in analyzing metabolic heterogeneity is the propensity for metabolite levels and labeling to change during cell handling and purification [39–41]. To evaluate this, we collected the single cell suspension right before MACS and compared metabolomics and labeling to directly- quenched tumor and spleen specimens. The purification process substantially altered the concentrations of most metabolites, including serine and glycine (Fig. S2C–D). It also diluted their labeling (Fig. S2E–F), reflecting their production, likely from protein catabolism during the purification process (further evidenced by increased leucine and isoleucine, Fig. S2C). ATP labeling, however, was largely preserved (Fig. 2B). This reflects ATP being a high abundance metabolite that cannot be readily produced during the nutrient-poor conditions of cell isolation. Notably, ATP catabolism will not alter the labeling pattern of the residual ATP. Thus, we were able to use ATP labeling from sorted cells to identify the extent of *de novo* versus salvage synthesis as the source of purines in TILs versus splenic T cells.

Purine sources were strikingly different between CD8^+^ TILs and splenic T cells: The splenic T cells, like spleen, had much higher fraction of purine labeling from inosine (salvage) than serine (*de novo*). Conversely, the TILs had much higher purine labeling from serine than inosine. With the purine fraction from inosine remaining similar, the upregulated purine fraction from serine in CD8^+^ TILs distinguished them from splenic T cells (Fig. 2C–D). Thus, CD8^+^ T cells within tumors stand out for high *de novo* purine synthesis.

### Both serine and circulating formate contribute to tumor 1C units

Having established a high *de novo* contribution to purines in tumors and TILs, we next sought to understand the sources of the 1C units supporting purine synthesis in these contexts (Fig. 3 A). Two important ones are serine’s hydroxymethyl carbon and formate. Accordingly, we infused MC38 tumor-bearing mice with [U-^13^C]serine or [^13^C]formate for 12 h and measured ATP labeling. Tumor ATP was labeled substantially by both tracers (Fig. 3B).

**Fig. 3.**
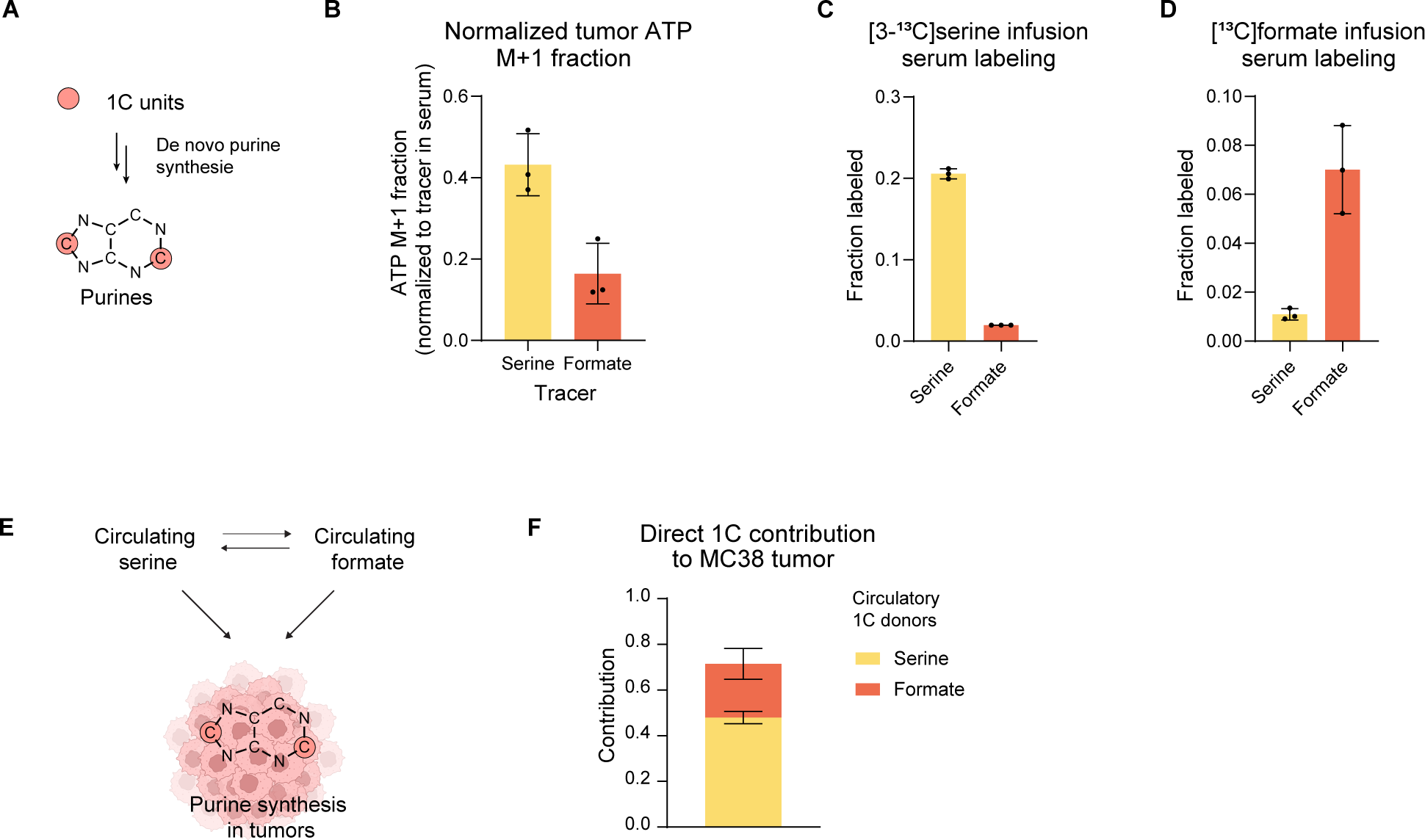
MC38 tumors synthesize purines *de novo* with 1C units from serine and circulating formate. (A) Schematic of 1C unit incorporation into the purine ring via *de novo* purine synthesis. (B) ATP M+1 labeled fraction in MC38 tumor, normalized to serum tracer enrichment for [U-^13^C]serine and [^13^C]formate infusions (n = 3). (C–D) Circulating serine and formate labeling in (C) [3-^13^C]serine or (D) [^13^C]formate infusions (n = 3). (E) Schematic representation of the potential for serine and formate to interconvert or directly feed tumor 1C units. (F) Direct contribution of circulating serine and formate to 1C units in MC38 tumor, determined by linear algebra with data from (B–D). Mean ± SD in (B–D). Mean ± SEM in (F). n indicates the number of mice.

Serine was only slightly labeled from formate (Fig. 3C), and vice versa (Fig. 3D), implying that most circulating formate comes from a source other than serine, perhaps microbiome metabolism. The observed limited cross-labeling between serine and formate suggest that both of their contributions to tumor ATP are largely direct. To quantify these contributions, we recognized that the steady state ATP M+1 fraction, *L*_*ATM,M+1*_, is related to 1C unit labeling (*L_1C_*) by

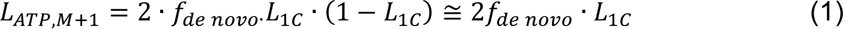

in which *f*_*de novo*_is the fraction of ATP from *de novo* synthesis (measured independently in Fig. 1H). We further corrected by linear algebra for any cross-labeling (Fig. 3E) [42], to arrive at a direct 1C contribution from each tracer, which was about 2x larger for circulating serine than for formate, with the combination accounting for 75% of all tumor purine 1C units (Fig. 3F). From the serine tracer, tumor serine was less labeled than circulating serine, presumably reflecting *de novo* serine synthesis within the tumor (Fig. S3A). Correcting also for this serine led to nearly complete accounting for tumor 1C units, roughly 80% from serine and 20% from formate (Fig. S3B).

We then carried out similar analysis of purine 1C unit sources from CD8^+^ TILs isolated from these tumors. They also showed a primary contribution from serine with a meaningful additional contribution from formate (Fig. S3C). Thus, both tumors as a whole and TILs have upregulated *de novo* purine synthesis supported by 1C units coming from serine and circulating formate.

### Increasing circulating formate drives its usage by T cells

We hypothesized that formate’s contribution to CD8^+^ TIL 1C units is limited by its circulating level (50 µM in mouse serum). To explore this, we carried out perturbative [^13^C]formate infusions, increasing total serum formate concentrations by up to 6 fold (Fig. 4A). This led to increased labeling in both CD8^+^ TILs (Fig. 4B) and splenic CD8^+^ T cells (Fig. 4C). Correcting for the labeled formate fraction in the circulation, the contribution of formate to total purine 1C units increased preferentially in the TILs (Fig. 4D). Circulating serine labeling was also modestly elevated, but much less than formate or the 1C unit enrichment in CD8^+^ TILs (Fig. S4A). Thus, increasing formate availability drives its direct usage in TIL purine synthesis.

**Fig. 4.**
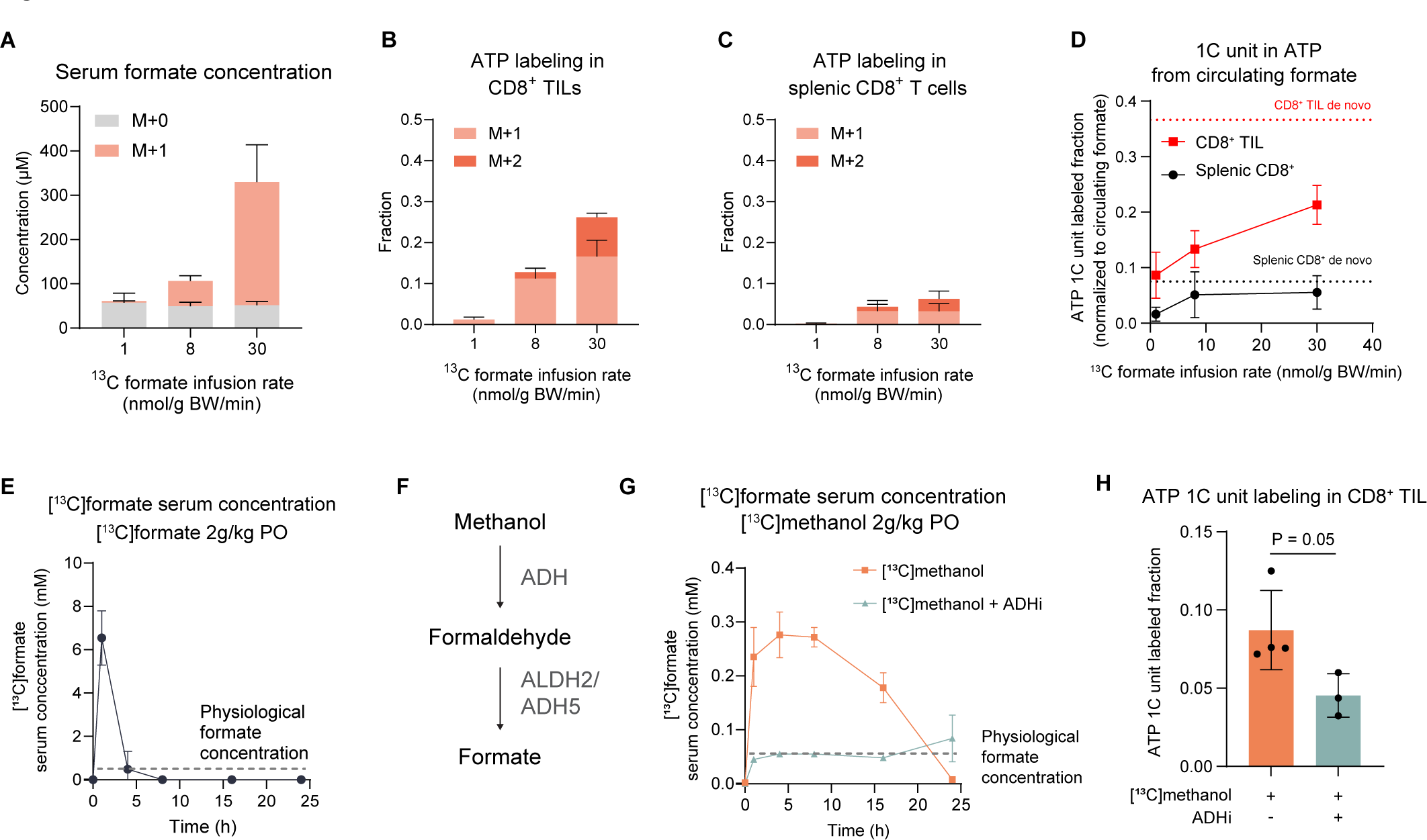
Experimental augmentation of circulating formate drives its usage by CD8^+^ TILs. (A) Circulating formate concentration and labeling achieved by infusing [^13^C]formate at different rates (n = 3). (B) Resulting ATP labeling in CD8^+^ TILs (n = 4 for 8 nmol/g BW/min; n = 3 for others). (C) Resulting ATP labeling in splenic CD8^+^ T cells (n = 3 for 1 nmol/g BW/min; n = 5 for others). (D) Relationship between [^13^C]formate infusion rate and circulating formate contribution to total ATP 1C units in CD8^+^ TILs and splenic CD8^+^ T cells. Calculated from data in (A–C). Dashed lines: fractions of purine from *de novo* synthesis (Fig. 2C). (E) Serum [^13^C]formate concentration over time after 2 g/kg [^13^C]formate oral gavage (PO) (n = 3). Dashed line: physiological formate concentration. (F) Schematic of methanol metabolism to formate. (G) Serum [^13^C]formate concentration over time after 2 g/kg [^13^C]methanol PO, with (n = 3) or without (n = 5) ADH inhibitor (fomepizole, 200 mg/kg PO). Dashed line: physiological formate concentration. (H) Labeling of ATP 1C units in CD8^+^ TILs 12 h after 2 g/kg [^13^C]methanol PO with and without ADH inhibitor (fomepizole, 200 mg/kg PO) (n = 4 for methanol alone; n = 3 for methanol with fomepizole). Mean ± SD in (E) and (G). Mean ± SEM in others. n indicates the number of mice.

As augmentation of CD8^+^ TIL purine synthesis could be beneficial for anti-tumor immunity, we looked for a more convenient way to elevate circulating formate levels. Formate becomes toxic at concentrations in the range of 5–10 mM [43], with pathological formate elevations responsible for the ocular toxicity (blindness) resulting from uncontrolled methanol ingestion. Oral gavage administration of [^13^C]formate (2 g/kg) resulted in a transient increase in circulating formate, which reached potentially toxic levels at 1 h and declined to baseline within 5 h (Fig. 4E).

The liver can convert methanol into formate, with methanol itself at least as safe as ethanol, except for its potential to lead to toxic formate accumulation (Fig. 4F) [44]. Accordingly, we explored the possibility that oral methanol gavage could be used to produce more stable circulating formate elevations. [^13^C]methanol (2 g/kg) elevated formate several-fold for over 12 h, while staying more than 10-fold below the concerning formate level of 5 mM (Fig. 4G). Such labeled methanol administration was sufficient to contribute substantially to CD8^+^ TIL purine synthesis (Fig. 4H). When ADH is inhibited by fomepizole, both [^13^C]formate production and its incorporation into CD8^+^ TIL decreased (Fig. 4G–H). Thus, methanol is a formate prodrug that elevates circulating formate concentrations and provides 1C units for TILs.

### Methanol supplementation synergizes with checkpoint inhibition

Based on the hypothesis that T cells responding to checkpoint blockade can sometimes be limited by insufficient 1C units (Fig. 5A), we next explored the potential for methanol to augment the effectiveness of checkpoint blockade. Mice bearing 100 mm^3^ MC38 tumors received anti-PD1 antibody (or isotype control) on four occasions over 10 days, with or without daily gavage of methanol 2 g/kg (Fig. 5B). In the absence of anti-PD-1, methanol had no impact on tumor growth (Fig. S5A). In combination with checkpoint blockade, however, methanol augmented tumor growth suppression (Fig. 5C). This favorable interaction was strong in some experiments and not others, consistent with inherent variability of MC38 immunotherapy response (Fig. S5B–E). Importantly, integrating data from four independent replicates, each with at least 10 mice per treatment group, the combination of methanol and anti-PD-1 increased markedly the frequency of durable regressions (Fig. 5D). Consistent with the mechanism being via formate generation, supplementation of drinking water with formate showed a trend towards beneficial interaction with anti-PD-1 (Fig. S5F–G), with formate drinking water recently reported to show synergy with checkpoint blockade in another murine cancer model (B16-OVA) [45]. Consistent with *de novo* purine synthesis contributing more than salvage in TILs, we did not observe benefits from oral inosine (Fig. S5H–J). This agrees with previous findings in the same tumor model [46]. While prior literature showed benefits from inosine supplementation in a different model [34], these did not include durable regressions, the desired clinical outcome of immunotherapy. Regressions induced by anti-PD-1 + methanol were associated with persistent antitumor immunity, as reimplantation of fresh MC38 tumor cells resulted in tumor rejection in all mice that responded durably to their initial tumors (Fig. 5E, Fig S5K). Thus, 1C unit supplementation via methanol as a formate prodrug augments the activity of anti-PD-1 immunotherapy, increasing durable regressions.

**Fig. 5.**
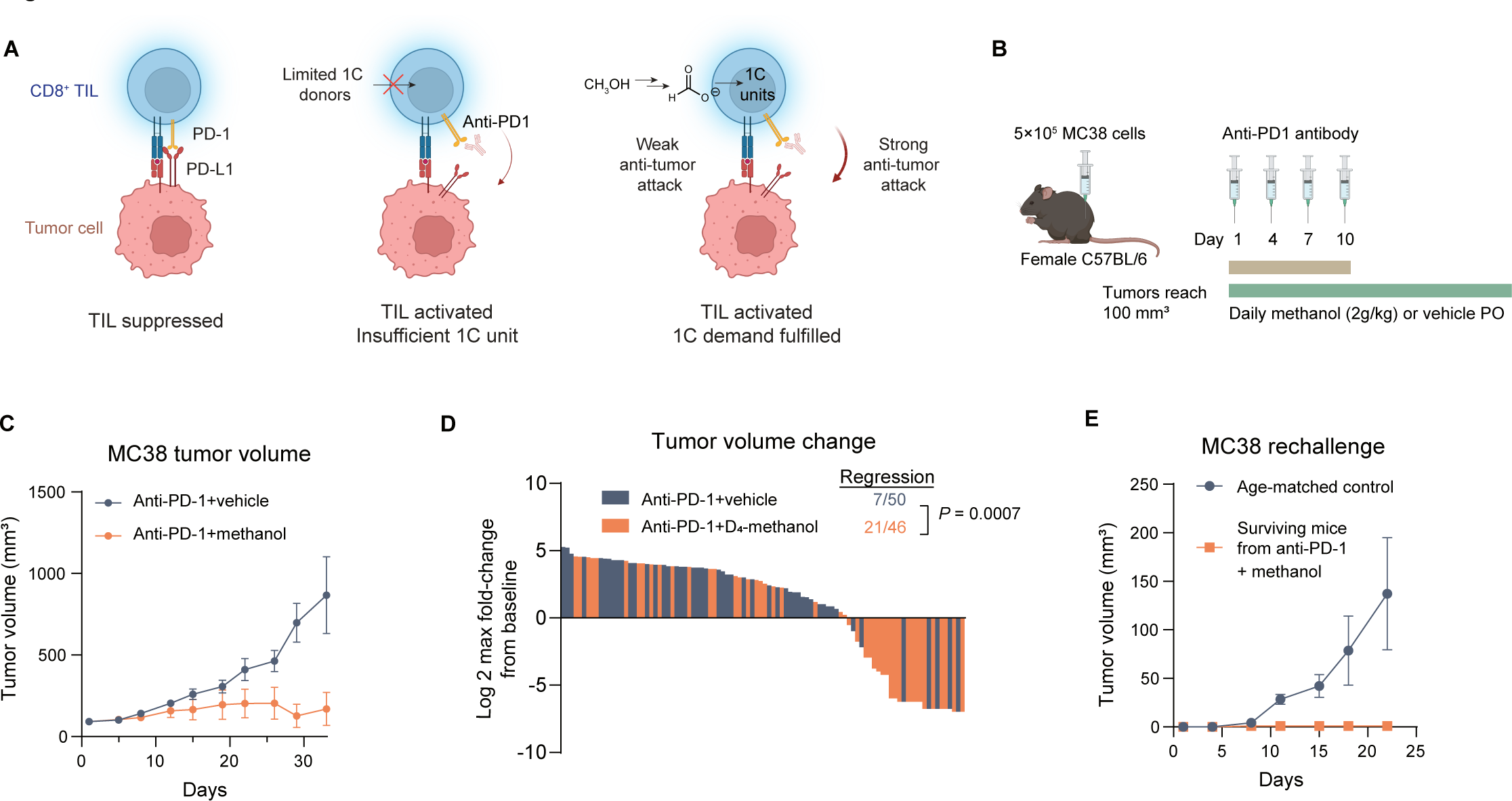
Methanol supplementation synergizes with PD-1 inhibition (A) Model for methanol supplementation boosting PD-1 inhibition. (B) Experimental design. vehicle = 0.9% sodium chloride in water. (C) MC38 tumor growth with anti-PD-1 ± methanol (n = 10). (D) Waterfall plot showing the maximal change in tumor volume compared to baseline (Day 1) for each mouse, pooling data from four independent experiments (Fig. S5B-E) with anti-PD-1 ± methanol. n is indicated on the plot. *P*-value was determined by Pearson chi-square test. (E) Tumor growth after MC38 rechallenge in surviving mice from anti-PD-1 + methanol treatment group (n = 6) and age-matched controls (n = 15). Mean ± SEM. n indicates the number of mice.

### Deuterium kinetic isotope effect can be used to control formate production rates

The primary safety risk of methanol ingestion is toxic formate buildup. This occurs when formate production by methanol oxidation outpaces formate clearance. In rodents, the balance of these reactions favors clearance of formate, rendering large doses of methanol safe, but mammals have lower formate clearance capacity [44]. We hypothesized that deuteration would slow methanol oxidation to formate, providing extended formate exposure at lower (safer) levels. In mice treated with [^13^C, D_4_]methanol, labeled plasma formate peaked around 150 μM, half the level of those treated with undeuterated [^13^C]methanol, and declined more gradually (Fig. 6A–C). Similar patterns were also observed in cynomolgus monkeys (Fig. 6B–C): Deuteration doubled the half-life of methanol (Fig. 6B), with 400 mg/kg steadily elevating plasma formate to 100 µM for 16 h (Fig. 6C). Clearance of formate itself was not affected by deuteration (Fig. S6A).

**Fig. 6.**
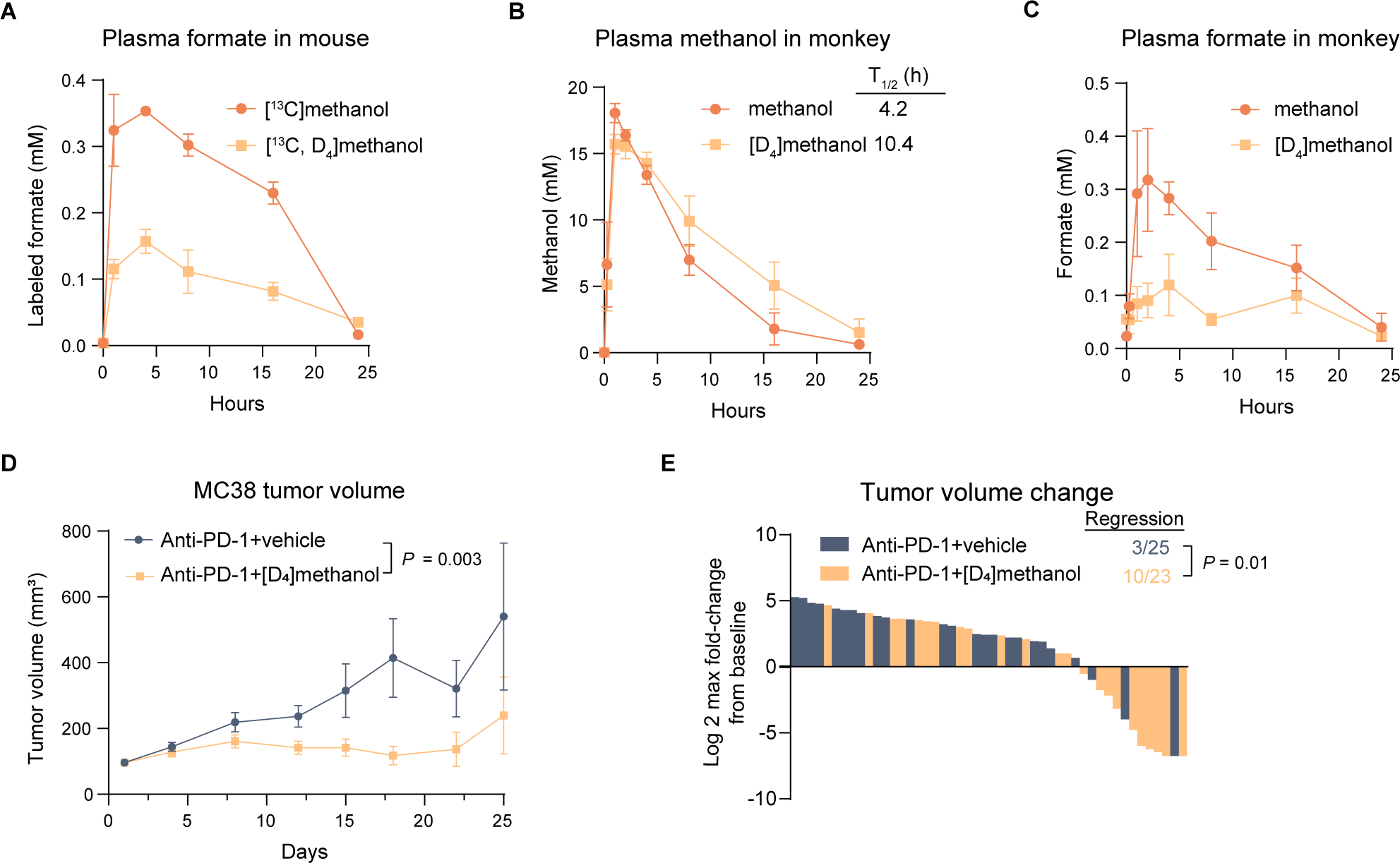
Deuterated methanol is a slow-release formate prodrug that augments immunotherapy (A) Plasma [^13^C] or [^13^C, D_4_]formate concentration over time after oral gavage of [^13^C] or [^13^C, D_4_]methanol at 2 g/kg in mice (n = 3). (B) Plasma methanol and (C) formate concentration over time after oral gavage of methanol or [D_4_]methanol at 400 mg/kg in monkeys (n = 3). (D) MC38 tumor growth with anti-PD-1 ± 2 g/kg [D_4_]methanol (n = 15 for vehicle, n = 13 for [D_4_]methanol). *P*-value was determined by a mixed-effects model for multiple comparisons with Geisser-Greenhouse correction. (E) Waterfall plot showing the maximal change in tumor volume compared to baseline (Day 1) for each mouse, pooling data from 2 independent experiments (Fig. S6B–C) with anti-PD-1 ± methanol. n is indicated on the plot. *P*-value was determined by Pearson chi-square test. Mean ± SD in (A–C). Mean ± SEM in (D). n indicates the number of animals.

We next examined whether [D_4_]methanol, like regular methanol, augments the activity of checkpoint blockade. Mice bearing MC38 tumors were treated with anti-PD-1 antibody, with and without daily gavage of [D_4_]methanol (2 g/kg). Despite subtler elevation of circulating formate (Fig. 6A), tumor suppression was equally strong (Fig. 6D), with the fraction of durable tumor regressions tripled across two independent experiments (Fig. 6E, Fig. S6B–C), mirroring the result with regular methanol at the same dose (Fig. 5D). Thus, deuterated methanol is a slow-release formate prodrug with the capacity to enhance immunotherapy.

## Discussion

Both cancer and activated immune cells reprogram metabolism to support their biosynthetic demands. One consistently upregulated pathway is 1C metabolism [19,47]. Its enzymes support both tumor and T cell expansion [23] and T cell differentiation into effector subsets [24]. A major function of 1C metabolism is to support *de novo* purine synthesis. Here, using stable isotope tracing, we show high activity of *de novo* purine synthesis in tumor-infiltrating T cells compared to splenic CD8^+^ T cells. The 1C unit supply supporting such synthesis is a potential metabolic immune checkpoint, and we show that supplementing 1C units systemically synergizes with PD- 1 inhibition.

The benefits of 1C supplementation for cancer immunotherapy stand in contrast to the importance of 1C enzyme inhibitors (antifolates) in cancer treatment. In current clinical oncology practice, the most important antifolate is pemetrexed, which preferentially targets thymidine synthesis. In contrast, methotrexate preferentially targets purine synthesis, and is now more commonly used as an immunosuppressant than cancer treatment. Indeed, low dose methotrexate to treat rheumatoid arthritis was shown to increase the risk of squamous cell carcinoma, likely due to immune suppression [48]. Thus, rather than considering the 1C pathway as a single entity, it is important to focus on specific enzymes, metabolites, and downstream products, with thymidine particularly important for cancer cells, and purines particularly important for immune cells.

To achieve 1C supplementation, we employed methanol as a formate prodrug. Methanol is converted by the sequential action of alcohol dehydrogenase and aldehyde dehydrogenase to formate, which is assimilated in the cytosol by the enzyme MTHFD1 to make 10-formyl-THF, a key purine biosynthetic input. Such supplementation did not just slow tumor growth under PD-1 inhibition, but resulted in durable tumor regressions in many mice, as did formate in the drinking water in a different tumor model in a recent report [45]. This differs from most metabolic strategies combined with anti-PD-1 to date, which tend to slow tumor growth but not result in durable regressions, e.g. supplementation of inosine [34], methionine [12], and arginine [49]. Instead, the preclinical results are more similar to, although somewhat less strong than, combination checkpoint therapies such as PD-1 and CTLA-4 [50,51], which are clinically effective [52,53].

From a safety perspective, methanol is well known for its toxic effects on the visual system, causing blindness when consumed erroneously instead of ethanol. These toxic effects are not caused by methanol directly, but by formate, when it accumulates in excess of 5 mM [43]. At such high concentrations, formate results in metabolic acidosis and respiratory chain impairment, damaging the retina. The immunotherapy-enhancing formate concentrations achieved here are up to 50x lower, and can be maintained steadily at this level in monkeys using [D_4_]methanol.

Interestingly, in human cancer patients, formate is among the most severely depleted circulating metabolites [54]. Moreover, folates, which are required for formate assimilation, are of lower abundance in humans than in mice [55]. Thus, it is tempting to speculate that 1C insufficiency is a yet bigger barrier to immunotherapy efficacy in the clinic, than it is in mouse models. Clinical studies of 1C supplementation are merited to address this possibility.

## Methods

### Experimental model and subject details

All mouse experiments were approved by the Institutional Animal Care and Use Committee at Princeton University and Charles River Laboratories. Experiments for testing anti-PD-1 efficacy were performed at Charles River Laboratories. All other mouse experiments were performed at Princeton University. Unless otherwise noted, C57BL/6 mice from Charles River Laboratories were used at the age of 9 to 15 weeks. Mice were housed under a normal light cycle (7 AM to 7 PM), with water and food (PicoLab Rodent Diet 5053, LabDiet) provided ad libitum. Jugular vein and/or carotid artery catheterization was performed in house. With sterile technique, catheters were inserted into the right jugular vein and (for arterial sampling) the left carotid artery and connected to a vascular access button implanted subcutaneously on the back of the mouse. Mice were allowed to recover for at least 5 days after surgery. To maintain catheter patency, catheters were flushed weekly with heparin saline (10 IU/mL) and locked by a catheter locking solution (500 IU/mL heparin in 50% glycerol, SAI Infusion Technologies).

Experiments involving cynomolgus monkeys (Macaca fascicularis) were approved by the Institutional Animal Care and Use Committee at WuXi AppTec Co.

### Mouse blood sampling

Blood was collected from the arterial catheter or by tail bleed. For arterial sampling, a cannula was connected to the vascular access button and mice were allowed to recover from handling-induced stress for at least 30 min. Blood was collected from the cannula directly to a collection tube without handling the mouse. For blood collection in the fasted state, mice were transferred to new cages without food at 9 AM, and blood was collected from 3 PM to 6 PM. Blood was kept on ice for up to 1 h followed by centrifugation (10,000 × g for 3 min, room temperature) as indicated. Serum was isolated from the supernatant and was extracted by methanol immediately (2.5 µL of serum into 70 µL of ice-cold methanol) or stored in -80°C until further analysis.

### Measurement of circulatory turnover flux for nucleosides

All measurements were performed in the fasted state. Mice were transferred to new cages without food at 9 AM. Water was supplied as HydroGels (ClearH_2_O). Starting at 2 PM, uniformly ^15^N- labeled nucleosides (dissolved in 0.9 % NaCl) were infused into the jugular vein. For efficiency, one purine tracer and one pyrimidine tracer were infused simultaneously into each mouse. Infusion parameters were summarized in Supplementary Table 1. Blood was sampled from the arterial catheter 120 min and 150 min after the start of infusion, and by tail bleed after the second arterial sampling. Blood was kept on ice after sampling. For measurement of purines, serum was isolated within 10 min from blood collection by centrifugation (10,000 × g for 3 min) and extracted by methanol (2.5 µL of serum into 70 µL of ice-cold methanol).

### Cell lines

The MC38 cell line was obtained from Kerafast (Boston, MA) and maintained at 37°C and 5% CO_2_ in Dulbecco’s Modified Eagle Medium (10-017-CV, Corning) supplemented with 10 vol% fetal bovine serum. To initiate tumors for infusion studies, cells were trypsinized, washed once with phosphate buffered saline (PBS), resuspended in PBS and kept on ice. 0.5 – 1 × 10^6^ cells were injected subcutaneously in 100 μL of PBS on the left flank of each mouse. Tumors were measured twice per week, with tumor size calculated by 0.5×length×width×height. Tracing experiments were performed when tumors reached 150-450 mm^3^ (2-3 weeks after implantation).

### Mouse infusion

Unless otherwise indicated, infusions were performed from 7 PM to 8 AM of the following day to trace purine ring turnover. Mice were single-housed with food and water provided *ad libitum*. Infusion parameters are summarized in Supplementary Table 1. At the end of infusion, blood was collected by tail bleed, and mice were euthanized by cervical dislocation. Spleens and tumors were cut in half, with one half for cell sorting and the other half for snap-freezing. To snap-freeze, tissues were wrapped in foils, clamped by a Wollenberger clamp precooled in liquid N_2_, and dropped in liquid N_2_.

### Formate and methanol dosing in monkeys

Restrained, non-sedated adult male monkeys were fasted overnight and re-fed 4 hours after dosing. Methanol or sodium formate was dissolved into saline and filtered through a 0.22 μm filter prior to administration. Intravenous doses were delivered as a bolus injection into a peripheral vein, while oral doses were delivered via nasogastric tube. At the indicated times, approximately 0.5 mL of whole blood was collected into potassium-EDTA tubes and rapidly processed into plasma, which was stored at -80°C until analysis.

### Magnetic activated cell sorting and metabolite extraction

Procedures were adapted from [10]. Spleens and tumors were cut into halves, and one half was used for cell sorting. Single-cell suspensions of splenocytes were prepared by manual disruption and passage through 70 µm cell strainers into PBS. Tumors were chopped and digested in 5 mL of PBS with 435 U/mL DNase (Sigma-Aldrich, D5025) and 218 U/mL collagenase (Sigma-Aldrich, C2674) at 37°C for up to 1 h on a shaker set at 200 rpm. The digested tumor cells were passed through 70 µm cell strainers into PBS. The splenocyte and tumor cell suspensions were then fractionated using CD8 (TIL) microbeads (Miltenyi Biotec, 130-116-478) according to the manufacturer’s instructions. Briefly, cells were resuspended in 10 µL of microbeads and 90 µL of MACS buffer (0.5% BSA + 2 mM EDTA in PBS) per 10^7^ cells at 4°C for 15 min. The magnetically labeled cells were loaded on LS columns (Miltenyi Biotec, 130-042-401) which were secured on Miltenyi MidiMACS Separators. Columns were washed twice, each time with 3 mL of MACS buffer to allow unlabeled cells to pass (collected as the CD8^-^ fraction). Columns were then removed from the MidiMACS Separators and flushed with 3 mL MACS buffer to collect the CD8^+^ fraction. The fractionated cells were resuspended in 1 mL of MACS buffer, with 20 µL used for cell counting, 900 µL for metabolic extraction, and all the remaining (∼80 µL) stained for flow cytometry.

To extract metabolites, cells were mixed with ice-cold acetonitrile:methanol:water (2:2:1, supplemented with 0.5 vol% formic acid which increases triphosphate yield [56]). 20 µL of extraction buffer per 10^6^ cells, or a total of 70 µL, whichever is more, was used. Extracts were vortexed for 10 s, and neutralized by NH_4_HCO_3_ (15% in water, 8.8% vol/vol of extraction buffer was used). Neutralized extracts were vortexed for 10 s again, kept on dry ice for 1h, and centrifuged (19,930 × g, 30 min at 4°C). Supernatants were collected and kept at -80°C until analysis.

### Flow cytometry analysis

Single-cell suspensions from tumors or spleens were incubated in Fc block (BD Biosciences, 553142, vol/vol 1/50) at room temperature for 10 min, washed once with staining buffer (PBS + 2% FBS), and stained for the following surface markers on ice for 30 min: CD4 (APC-Cy7, 1:100, clone RM4-5, BD Biosciences, 565650), CD8b (Alexa Fluor 488, 1:250, clone YTS156.7.7, Biolegend, 126628), CD45.2 (eFluor 450, 1:100, clone 104, Thermo Fisher, 48-0454-82). Either Live/Dead Aqua (Thermo Fisher, L34966) or propidium iodide (Thermo Fisher, R37169) was used as viability dye according to the manufacturer’s instructions. All flow cytometry was analyzed with an LSR II flow cytometer (BD Biosciences) and FCS Express 7.12 (De Novo Software). Gating strategies are shown in Supplementary Fig. 2A–B.

### Tissue metabolite extraction

Eppendorf tubes and ceramic beads were precooled on dry ice. Tissues were transferred to Eppendorf tubes and disrupted by cryomill (Retsch). About 10 mg of homogenized tissue powder was weighed and extracted by ice-cold acetonitrile:methanol:water (2:2:1, supplemented with 0.5 vol% formic acid). Extracts were vortexed for 10 s, and neutralized by NH_4_HCO_3_ (15% in water, 8.8% vol/vol of extraction buffer was used). Neutralized extracts were vortexed for 10 s again, kept on dry ice for 1h, and centrifuged (19,930 × g, 30 min at 4°C). Supernatants were collected and kept at -80°C until analysis.

### Formate derivatization

Analysis of formate is challenging due to unlabeled formate in procedure blanks, and this value was subtracted from the observed levels in experimental samples. Given the importance of this measurement, both GC-MS and LC-MS were used.

To measure [^13^C]formate concentration only (not unlabeled formate) in serum samples, an LC-MS-compatible derivatization method for carboxylic acids (adapted from [57]) was used. It enabled amide bond formation between carboxylic acid and 3-nitrophenylhydrazine (3-NPH) with coupling reagent 1-ethyl-3(3-(dimethylamino)propyl)carbodiimide (EDC). 2 µL of serum was added to 40 µL of derivatization reagent (25 mM 3-NPH, 12 mM EDC, 2.4 vol% pyridine, in methanol) and was incubated on ice for 1 h. Samples were spun (19,930 × g, 20 min at 4°C), and the reaction was quenched by adding 10 µL of supernatant to 90 µL of sodium propionate (5 mM in water). The mixture was incubated on ice for 10 min and centrifuged (19,930 × g, 10 min at 4°C) again. Supernatants were collected and kept at 4°C until analysis. [^13^C]formate concentration was extrapolated from standard curves after natural abundance correction.

To measure both unlabeled and ^13^C-labeled formate in serum samples, a GC-MS-compatible derivatization method (adapted from [36]) was used. In short, formate was converted to benzyl formate by reaction with benzyl alcohol and chloromethyl chloroformate (CMCF). A reaction mixture was prepared by adding 5 µL of NaOH (1 M, in water), 2.5 µL of benzyl alcohol, and 20 µL of pyridine into an Eppendorf tube. Then 10 µL of serum was added and mixed by pipetting. To initiate the reaction, 5 µL of CMCF was added, and tubes were quickly capped and vortexed for 15 s. The procedure must be performed in fume hoods as gaseous HCl is generated. The mixture was allowed to sit at room temperature for 20 min and then diluted by 100 µL of water and 100 µL of methyl t-butyl ether (MTBE). Samples were vortexed for 5 s, spun (19,930 × g, 10 min at 4°C), and the organic layer was transferred to glass vials. Vials were kept at room temperature until analysis. By GC-MS, a trace amount of benzyl formate was observed in benzyl alcohol (from various vendors) directly diluted in MTBE, which is likely an impurity from the reagent and contributes to the formate background in procedure blanks (∼30 µM). To subtract the background, standard curves with at least three procedure blanks were used to extrapolate formate concentration after natural isotope correction.

### LC-MS method

Metabolites were separated by hydrophilic interaction liquid chromatography (HILIC) with an XBridge BEH Amide column (2.1 mm × 150 mm, 2.5 μm particle size; Waters, 196006724). The column temperature was set at 25°C. Solvent A was 95 vol% H_2_O 5 vol% acetonitrile (with 20 mM ammonium acetate, 20 mM ammonium hydroxide, pH 9.4). Solvent B was acetonitrile. Flow rate was 0.15 mL/min. The LC gradient was: 0-2min, 90% B; 3-7min, 75% B; 8-9 min, 70% B; 10-12 min, 50% B; 12-14 min, 25% B; 16-20.5 min, 0.5% B; 21-25 min, 90%.

MS analysis was performed on Thermo Fisher’s Q Exactive Plus (QE+) Hybrid Quadrupole-Orbitrap, Orbitrap Exploris 240, or Orbitrap Exploris 480 mass spectrometer. Full scan was performed in negative mode, at the m/z of 70-1000. The automatic gain control (AGC) target was 3e6 on QE+ and 1000% on Exploris 240/480. The maximum injection time was 500 ms on QE+ and Exploris 480, and 200 ms on Exploris 240. The orbitrap resolution was 140,000 on QE+, and 180,000 on Exploris 240/480. Selected ion monitoring (SIM) was performed for ATP in negative mode at the retention time of 13-15min, m/z 505-515, AGC target 1e6 (QE+) or 1000% (Exploris 240/480), maximum injection time 500 ms, and orbitrap resolution 140,000 (QE+) or 240,000 (Exploris 240), or 480,000 (Exploris 480).

Derivatized formate (by reaction with 3-NPH) was analyzed by QE+. LC separation was done by a reversed-phase method, with an Acquity UPLC BEH C18 column (2.1 mm × 100 mm, 1.7 µm particle size, 130 Å pore size; Waters, 186002352) using a gradient of solvent A (water) and solvent B (methanol): 0-1min, 10% B; 5 min, 30% B; 7-10 min, 100% B; 10.5-18 min, 10% B. Flow rate was 0.2 mL/min. Selective ion monitoring was done in negative mode with m/z 178 to 186, AGC target 3e6, and maximum injection time 500 ms.

### GC-MS method

To analyze formate (derivatized to benzyl formate), 0.5 µL of sample was loaded on Agilent’s 7250 GC/Q-TOF in splitless injection mode. The injection port was set at 250°C. GC separation was performed on a VF-WAXms fused silica column (30 m × 0.25 mm, 0.5 μm film thickness, Agilent, CP9222), with the oven temperature programmed as follows: initiation at 56°C and hold for 3 min, then ramped to 220°C (ramp rate 30 °C/min) and hold for 5 min. Helium was used as the carrier gas at a constant flow rate of 1.1 mL/min. Electron ionization was set at the low energy mode (15 eV), and full scan was performed at m/z 50-500.

To analyze methanol, 50 µL of plasma was mixed with 150 µL of N-methyl-2-pyrrolidone containing acetonitrile as an internal standard (final concentration = 26 µg/mL) in glass vials. Vials were sampled with an Agilent 7697A Headspace Sampler and analyzed by a 7890B GC system coupled to a 5977B mass spectrometer. The column was an Agilent DB-WAX Ultra Inert GC column (30 m × 0.25 mm × 0.25 µm) using helium as a mobile phase at a flow rat of 1 mL/min. The temperature gradient was as follows: 30°C, 5 minutes; ramp to 230°C at a rate of 120° C/minute; hold for 3 minutes. Additional GC parameters: vial equilibration, 5 minutes; solvent delay, 1.6 minutes; headspace oven temp, 60°C; headspace loop temp, 120°C; MS transfer line temp, 280°C. Analytes were ionized by electron ionization and detected as the molecular ions. Additional MS parameters: source temp, 230°C; quadrupole temp, 150°C. Concentrations were determined using external standard curves prepared in plasma from untreated monkeys.

### Formate measurement by ion chromatography

To measure formate in the plasma from monkeys, 75 µL of plasma was diluted in 150 µL of acetonitrile and centrifuged (14,000 × g, 10 minutes). 150 µL of supernatant was diluted into 300 µL of water, and 25 µL was injected into a Dionex ICS-5000+ IC system using an IonPac AS18 RFIC column (250 mm × 4 mm) with an IonPac AG11-HC RFIC guard column (250 mm × 4 mm). The mobile phase was water with potassium hydroxide generated by a Dionex EGC 500 KOH cartridge. The KOH gradient was: 0 to 15 minutes, 8 mM; 15 to 15.1 minutes, ramp to 50 mM; 15.1 to 20 minutes, 50 mM; 20 to 20.1 minutes, drop to 8 mM; 20.1 to 30 minutes 8 mM. Flow rate was 1 mL/minute. Column temperature was 30°C. Analytes were detected by electron capture. Concentrations were determined using external standard curves prepared in plasma from untreated monkeys.

### PD-1 efficacy with nutrient supplementation

Anti-tumor efficacy experiments were performed at Charles River Laboratories. Female C57BL/6 mice were used which were 8-12 weeks old at the date of tumor initiation. Water and Rodent NIH- 31M Auto chow (Ziegler Feed) was provided *ad libitum*. To initiate tumors, 5×10^5^ MC38 cells in 1 mL PBS were injected subcutaneously on the flank of each mouse. When the average tumor volume reached 80-120 mm^3^, mice were randomized into different groups and were treated with anti-PD-1 (5 mg/kg, clone RMP1-14, ichorbio IHC1132) or isotype control Rat IgG2a (5 mg/kg, clone 2A3, BioXcell BE0089) twice per week for two weeks. Nutrient supplementation started from the first dose of antibody treatment and lasted until experimental endpoint. To supplement formate, sodium formate (2% w/v) was provided in the drinking water. To supplement formate via methanol, 2 g/kg of methanol (25% v/v in saline, dosing volume 10 mL/kg) was administered via daily oral gavage. To supplement inosine, 0.9 g/kg of inosine (90 mg/mL suspension in PBS, dosing volume 10 mL/kg) was administered twice per day (12 hour intervals) via oral gavage. For MC38 rechallenge, 5×10^5^ MC38 cells were injected subcutaneously on the opposite flank of tumor-free mice from anti-PD-1 ± methanol treatment or age-matched female C57BL/6 control mice. No antibody or methanol was administered. Tumor volumes were measured twice per week.

### Data analysis

Raw mass spectrometry data were converted to .mzXML format by MSConvert (ProteoWizard). Pick-peaking was done on El Maven (v0.8.0, Elucidata). Natural isotope abundance was corrected by the ‘accucor’ package on R for all metabolites except ATP, whose ^13^C_2_ (-H^+^ 507.9952 m/z) isotopomer is partially resolved from ^18^O_1_ (-H^+^ 507.9927 m/z). To address this, an algorithm (optCorr, available at https://github.com/xxing9703/optCorr_script) was developed. It accounts for the contribution from unresolved or partially-resolved peaks to the observed peak top (e.g., the contribution from ^18^O_1_ to the ^13^C_2_ peak top) by simulating MS profiles. Natural abundance correction for ATP was done by optCorr.

Circulatory turnover flux (F_circ_) of nucleosides was determined by the same method as previously described [42]. F_circ_ was calculated by

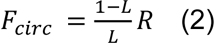

where L is the fraction of labeled nucleoside in arterial serum, and R is the rate at which such nucleoside was infused.

The fraction of newly synthesized ATP was calculated by the fractional purine ring carbon atom labeling from infusing [U-^13^C]serine. Such labeling comes from both labeled glycine and labeled 1C units. The resulting total ATP carbon atom labeling is given by ∑_*i*_ *i* × *L*_*ATP,M+i*_ with the M+3 and M+4 forms undetectable (as expected from the minimally perturbative serine infusion). The total possible labeling (if all ATP were new) is given by 2 × *L*_*gly,M+2*_ + 2 × (*L*_*Ser,M+1*_ + *L*_*ser,M+3*_). The newly synthesized ATP fraction is the ratio of these:

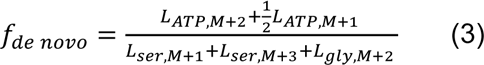

In isolated cells, serine and glycine labeling was diluted during sample preparation, so labeling measured in clamped bulk tissues was used instead.

The fraction of ATP with its purine ring from circulating inosine *f*_*insoinde*_ was calculated by normalizing the ATP labeling to serum allantoin as a surrogate for inosine.

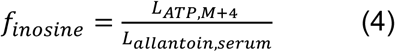

For [U-^15^N]adenosine infusion, tissue ATP labeling was normalized to arterial adenosine enrichment *L*_*adenosine,serum*_ measured independently in double catheterized mice.

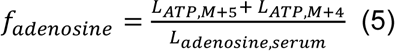

Note that the M+4 ATP (a relatively small fraction) can come from M+5 ATP via the purine nucleotide cycle or from circulating inosine labeled by adenosine. No correction was made to subtract adenosine->inosine->ATP contributions.

The direct contribution to the tumor 1C pool was determined by a linear algebra strategy previously described for other metabolites [42]. Both serine and formate from circulation can label the 1C pool for purine synthesis, and each can label the other. The goal is to figure out the direct 1C source correcting for the indirect routes (e.g. circulating formate -> circulating serine -> purine 1C). This is important for ensuring that there is a direct 1C contribution from circulating formate. In either [U-^13^C]serine or ^13^C-formate infusion, the fraction of labeled 1C pool in tissues *L*_1*C*_ can be accounted for by the sum of direct contribution of circulating serine and formate (*f*_*ser*_ and *f*_*for*_), and serine/formate labeling. For serine, both M+1 and M+3 can contribute to labeled 1C units, so we call *L*_*serum,ser*_1*C*__ = *L*_*serum,ser, M + 1*_ + *L*_*serum,ser, M + 3*_.

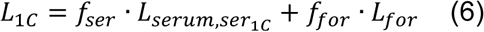

In [U-^13^C]serine infusion, define *L*_*ser→for*_ as the fraction of circulating formate from circulating serine:

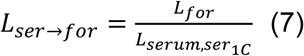

Thus, (6) becomes

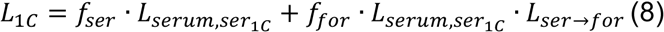

Similarly, in ^13^C-formate infusion,

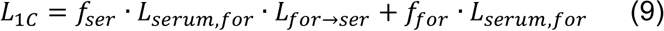

Substituting (8) and (9) into (1), and rearranging into matrix form gives:

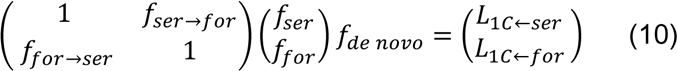

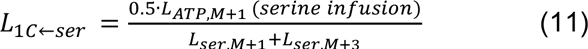

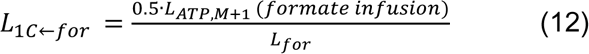

*f*_*ser*_ and *f*_*for*_ were calculated by fitting experimental data with (10-12), and errors were determined by bootstrapping.

To account for tumor serine which is not derived from circulation (e.g., *de novo* synthesis), we first calculated its fraction *L*_*ser,non−circ*_ by comparing the average carbon-atom labeling for serine in tumors and serum.

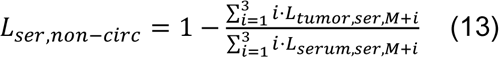

Its contribution to tumor’s 1C pool *f*_*ser,non−circ*_ was calculated by

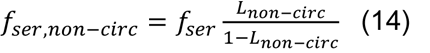

And errors were propagated by standard error propagation methods.

To determine the 1C pool labeling in isolated cells from perturbative ^13^C-formate infusions, we used the ratio of M+1 and M+2 fractions for ATP. The fraction of M+2 ATP is given by

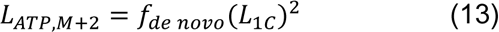

Solving for *L*_1*C*_ from (1) and (13) gives

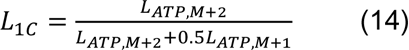

**Supplementary Table 1.**
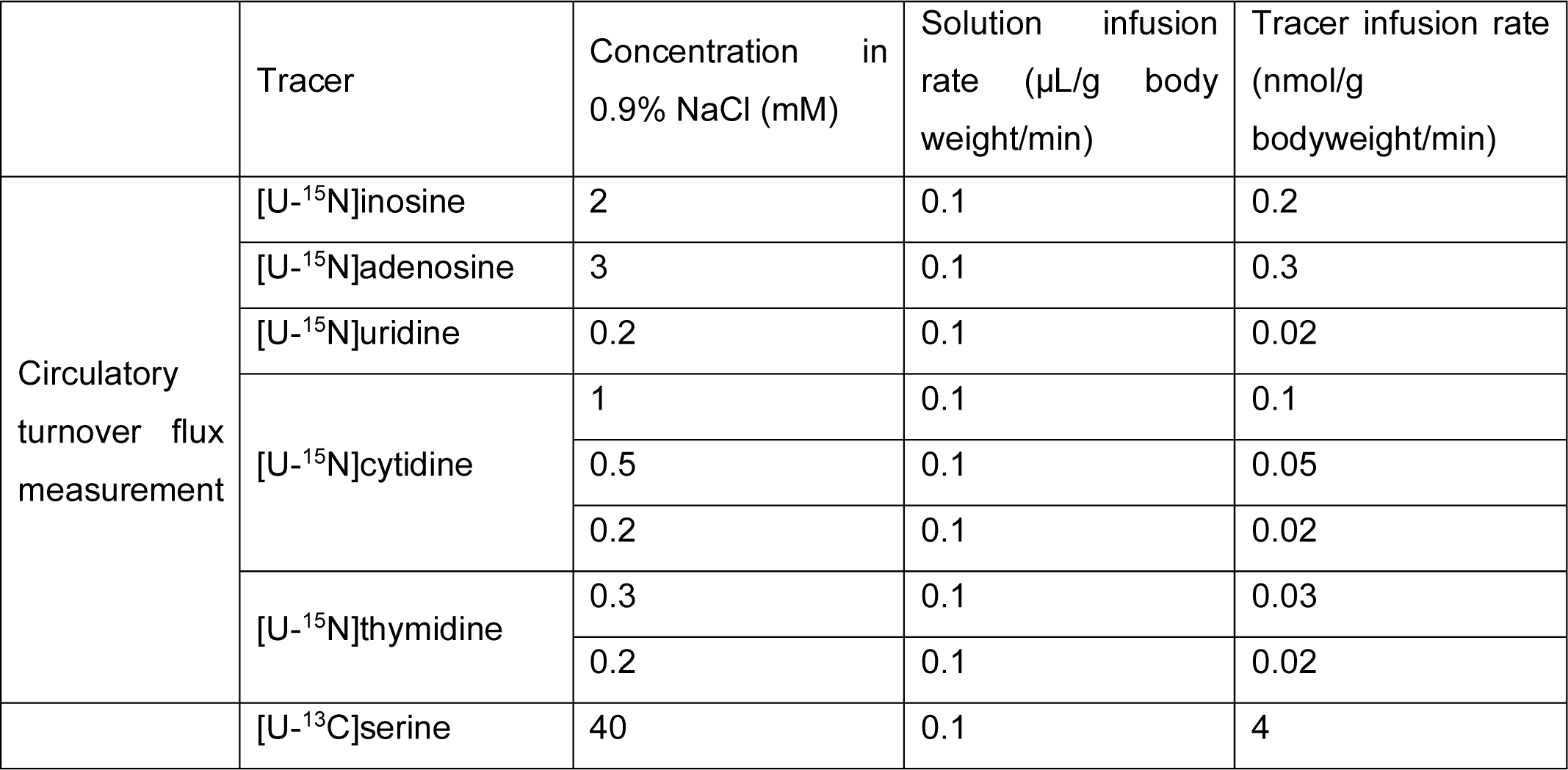
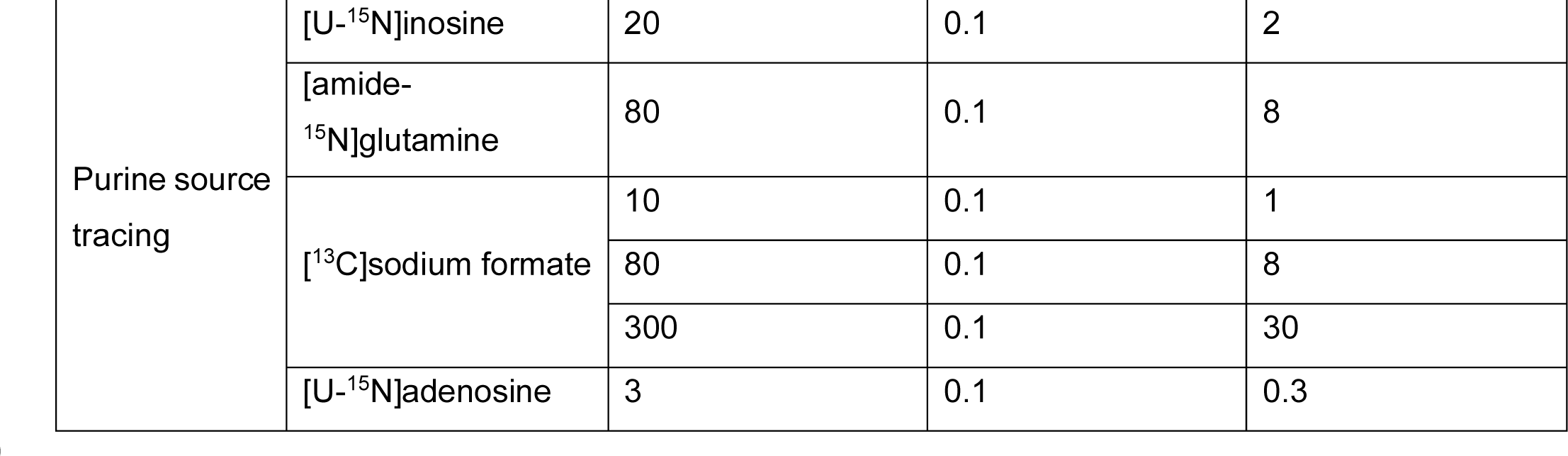
Infusion Parameters.

## Acknowledgements

J.D.R. was supported by Ludwig Cancer Research, R01CA163591 from the National Cancer Institute, and DP1DK113643 from the National Institute of Diabetes, Digestion and Kidney Disease. X.Xu and Z.C. were supported by the Edward C. Taylor Third Year Fellowship in Chemistry. C.R.B. was supported by the Damon Runyon Postdoctoral Fellowship. We thank John Eng at the Mass Spectrometry Core Facility for assistance with GC-MS and Christina DeCoste at the Flow Cytometry Resource Facility. We thank Wenyun Lu, Yihui Shen, Tara TeSlaa, Won Dong Lee, and all other Rabinowitz lab members for insights and discussion.

## Author contribution

J.D.R. and K.O. conceived the study. X.Xu and Z.C. came up with the general experimental strategy for in vivo tracing and performed most experiments. K.O. came up with the strategy for supplementing formate via methanol, designed and oversaw all PD-1 efficacy studies. Z.C. and C.R.B. optimized the cell isolation protocol. X.Xing wrote optCorr for natural isotope abundance correction. J.D.R., X.Xu, and Z.C. wrote the paper with inputs from all authors.

## Legends for Supplementary Figures

Supplementary Fig. 1. Complexity of measuring circulating nucleoside abundance and turnover.

(A) Certain purines are enriched in blood collected from mouse tail versus carotid artery catheter, reflecting production within the tail or during tail sampling. Serum was isolated and extracted within 10 min after blood collection (n = 6). *P*-values were calculated by unpaired two-tailed Student’s *t*-test.

(B) Volcano plot showing the change in serum metabolite levels when letting the blood sit on ice prior to serum isolation via centrifugation (comparing the minimum clotting time of 10 min versus a typical time of 30 min, n = 3).

(C) ^15^N nucleosides’ labeling is diluted in tail serum compared to artery, especially for inosine. n

= 3 for inosine and thymidine. n = 2 for uridine, adenosine, and cytidine. *P*-values were calculated by unpaired two-tailed Student’s *t*-test.

(D) Serum inosine and allantoin labeling in [U-^15^N]inosine infusion (n = 3). Allantoin sampling from either jugular artery or the tail vein accurately reflects arterial inosine labeling. Serum was processed within 10 min from blood collection. *P*-values were calculated by unpaired two-tailed Student’s *t*-test.

(E) Serum allantoin concentration is stable after blood collection (n = 3). *P*-values were calculated by unpaired two-tailed Student’s *t*-test.

(F–G) Labeling of inosine and adenosine in quickly processed arterial serum during (F) [U- ^15^N]inosine and (G) [U-^15^N]adenosine infusions (n = 4). Infused labeled adenosine is converted into labeled circulating inosine, but not vice versa.

(H) Kinetics of tissue ATP labeling during [amide-^15^N]glutamine infusion. n = 3 for spleen at 4 h, and n = 2 for others at each time point.

(I–K) Comparison of labeling in different purine nucleotides. In each graph, data points are from the same infusion, for different sampled tissues and measured nucleotides.

(I) Adenosine and guanosine nucleotides versus IMP from [U-^13^C]serine infusion (n = 3). All nucleotides label to the same extent.

(J) Adenosine and guanosine nucleotides versus IMP from [U-^15^N]inosine infusion (n = 3). Most nucleotides label to the same extent, with some trend towards greater labeling in IMP than downstream nucleotides.

(K) Adenosine, guanosine, and inosine nucleotides versus AMP from [U-^15^N]adenosine infusion (n = 3). Guanosine nucleotides label less than adenosine nucleotides in some tissues.

(L) Tissue ATP M+5 fraction (directly from adenosine) and M+4 fraction (from M+5 adenosine becoming inosine or IMP) in [U-^15^N]adenosine infusion (n = 3). Direct adenosine assimilation predominates.

Mean ± SD; n indicates the number of mice. ns, p > 0.05; * p < 0.05; ** p < 0.01; *** p < 0.001; **** p < 0.001.

Supplementary Fig. 2. Cell isolation yielded populations of good purity but altered the abundance and labeling of some metabolites.

(A–B) Representative flow cytometry result of all cells, CD8^+^, and CD8^-^ fractions in (A) MC38 tumor and (B) spleen gated on live CD45^+^ cells.

(C–F) Changes in metabolite abundance and labeling in dissociated MC38 tumor and spleen compared to the same snap-frozen sample (n = 3).

(C–D) Volcano plots showing changes in metabolite abundance. Metabolite abundance was normalized to the mean of all metabolites (log2 ion count) for each sample.

(E–F) The change of serine (E) and glycine (F) labeling (from [U-^13^C]serine infusion) *P*-values determined by paired two-tailed Student’s *t*-tests. n indicates the number of mice. ns, p > 0.05; * p < 0.05; ** p < 0.01.

Supplementary Fig. 3. 1C unit input in MC38 tumor and CD8^+^ TILs.

(A) Serine labeling pattern in serum and MC38 tumor for [U-^13^C]serine infusion. Mean ± SD. n = 3 mice.

(B) Direct contribution to tumor 1C units from circulating formate, circulating serine, and serine *de novo* synthesized in tumor. Mean ± SEM.

(C) Direct contribution of circulating serine and formate to 1C units in bulk MC38 tumor and CD8^+^ TILs. Mean ± SEM.

Supplementary Fig. 4. Elevated circulating formate feeds CD8^+^ TILs directly.

(A) Serum serine, serum formate, and 1C unit in CD8^+^ TILs labeling at different [^13^C]formate infusion rates. TIL labeling markedly exceeds circulating serine labeling, implying that the contribution of formate is direct and not primarily via circulating serine. Mean ± SD. At each infusion rate, n = 4 for serine and n = 3 for others.

Supplementary Fig. 5. Effect of supplementing methanol, formate, or inosine on anti-PD-1 efficacy.

(A) MC38 tumor growth with isotype antibody (5 mg/kg IP, BIW for 2 weeks) ± methanol (2 g/kg daily PO). n = 10.

(B–E) Individual MC38 tumor volumes from 4 independent experiments with anti-PD-1 (5 mg/kg IP, BIW for 2 weeks) ± methanol (2 g/kg daily PO). n is indicated on each panel.

(F–G) MC38 tumor growth with anti-PD-1 (5 mg/kg IP, BIW for 2 weeks) ± sodium formate (2% in drinking water). n = 10.

(F) Individual tumor volumes.

(G) Waterfall plot showing the maximal fold-change of MC38 tumor volume compared to baseline (start of treatment) for each mouse. n = 10 for each group. *P*-value was determined by Pearson chi-square test.

(H–J) MC38 tumor growth with anti-PD-1 (5 mg/kg IP, BIW for 2 weeks) ± inosine (0.9 g/kg BID PO). n = 15.

(H) Average tumor volumes.

(I) Individual tumor volume.

(J) Waterfall plot showing the maximal fold-change of MC38 tumor volume compared to baseline (start of treatment) for each mouse. n = 15. *P*-value was determined by Pearson chi-square test.

(K) Growth of individual tumors after MC38 rechallenge in surviving mice from anti-PD-1 + methanol treatment group (n = 6) and age-matched controls (n = 15).

Mean ± SEM in (A) and (H). n indicates the number of mice.

Supplementary Fig. 6. Pharmacokinetics of deuterated formate and bioactivity of deuterated methanol in combination with checkpoint blockade.

(A) Plasma formate concentration over time in monkeys intravenously inject with [^13^C] or [D]formate at 50 mg/kg. n = 3.

(B–C) Individual MC38 tumor volumes. Mice were treated with 4 doses of anti-PD-1 with and without daily gavage of 2 g/kg [D_4_]methanol. n is indicated on each panel.

Mean ± SD. n indicates the number of animals.

