## Supplemental Figures for "One-carbon unit supplementation fuels tumor-infiltrating T cells and augments checkpoint blockade"

Figure S1

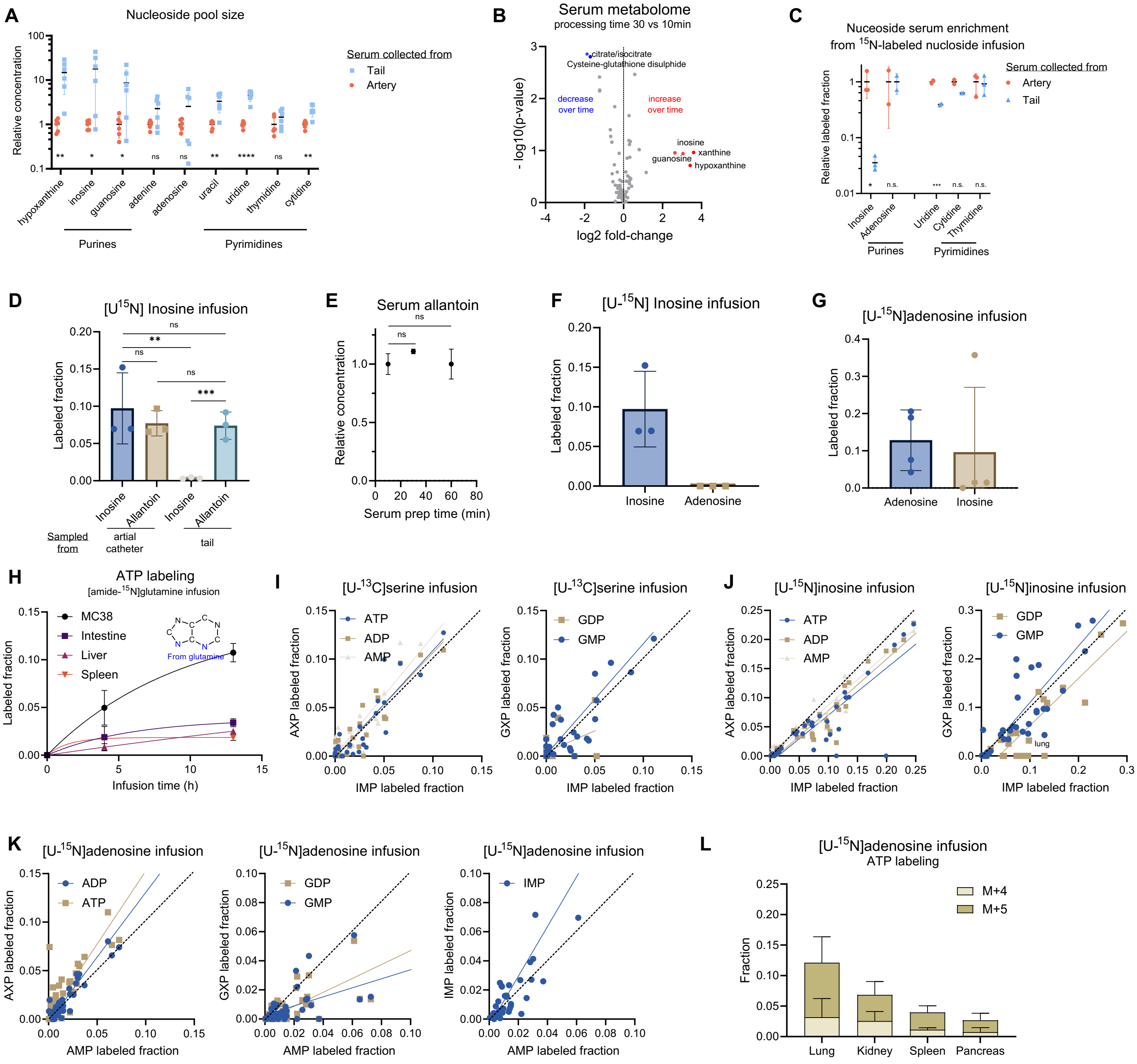


**Supplementary Fig. 1. Complexity of measuring circulating nucleoside abundance and turnover.**
(A) Certain purines are enriched in blood collected from mouse tail versus carotid artery catheter, reflecting production within the tail or during tail sampling. Serum was isolated and extracted within 10 min after blood collection (n = 6). *P*-values were calculated by unpaired two-tailed Student's *t*-test.
(B) Volcano plot showing the change in serum metabolite levels when letting the blood sit on ice prior to serum isolation via centrifugation (comparing the minimum clotting time of 10 min versus a typical time of 30 min, n = 3).
(C) $^{15}$N nucleosides' labeling is diluted in tail serum compared to artery, especially for inosine. n = 3 for inosine and thymidine. n = 2 for uridine, adenosine, and cytidine. *P*-values were calculated by unpaired two-tailed Student's *t*-test.
(D) Serum inosine and allantoin labeling in [U-$^{15}$N]inosine infusion (n = 3). Allantoin sampling from either jugular artery or the tail vein accurately reflects arterial inosine labeling. Serum was processed within 10 min from blood collection. *P*-values were calculated by unpaired two-tailed Student's *t*-test.
(E) Serum allantoin concentration is stable after blood collection (n = 3). *P*-values were calculated by unpaired two-tailed Student's *t*-test.
(F–G) Labeling of inosine and adenosine in quickly processed arterial serum during (F) [U-$^{15}$N]inosine and (G) [U-$^{15}$N]adenosine infusions (n = 4). Infused labeled adenosine is converted into labeled circulating inosine, but not vice versa.
(H) Kinetics of tissue ATP labeling during [amide-$^{15}$N]glutamine infusion. n = 3 for spleen at 4 h, and n = 2 for others at each time point.
(I–K) Comparison of labeling in different purine nucleotides. In each graph, data points are from the same infusion, for different sampled tissues and measured nucleotides.
(I) Adenosine and guanosine nucleotides versus IMP from [U-$^{13}$C]serine infusion (n = 3). All nucleotides label to the same extent.
(J) Adenosine and guanosine nucleotides versus IMP from [U-$^{15}$N]inosine infusion (n = 3). Most nucleotides label to the same extent, with some trend towards greater labeling in IMP than downstream nucleotides.
(K) Adenosine, guanosine, and inosine nucleotides versus AMP from [U-$^{15}$N]adenosine infusion (n = 3). Guanosine nucleotides label less than adenosine nucleotides in some tissues.
(L) Tissue ATP M+5 fraction (directly from adenosine) and M+4 fraction (from M+5 adenosine becoming inosine or IMP) in [U-$^{15}$N]adenosine infusion (n = 3). Direct adenosine assimilation predominates.
Mean ± SD; n indicates the number of mice. ns, $p > 0.05$; * $p < 0.05$; ** $p < 0.01$; *** $p < 0.001$; **** $p < 0.001$.

**Figure S2**

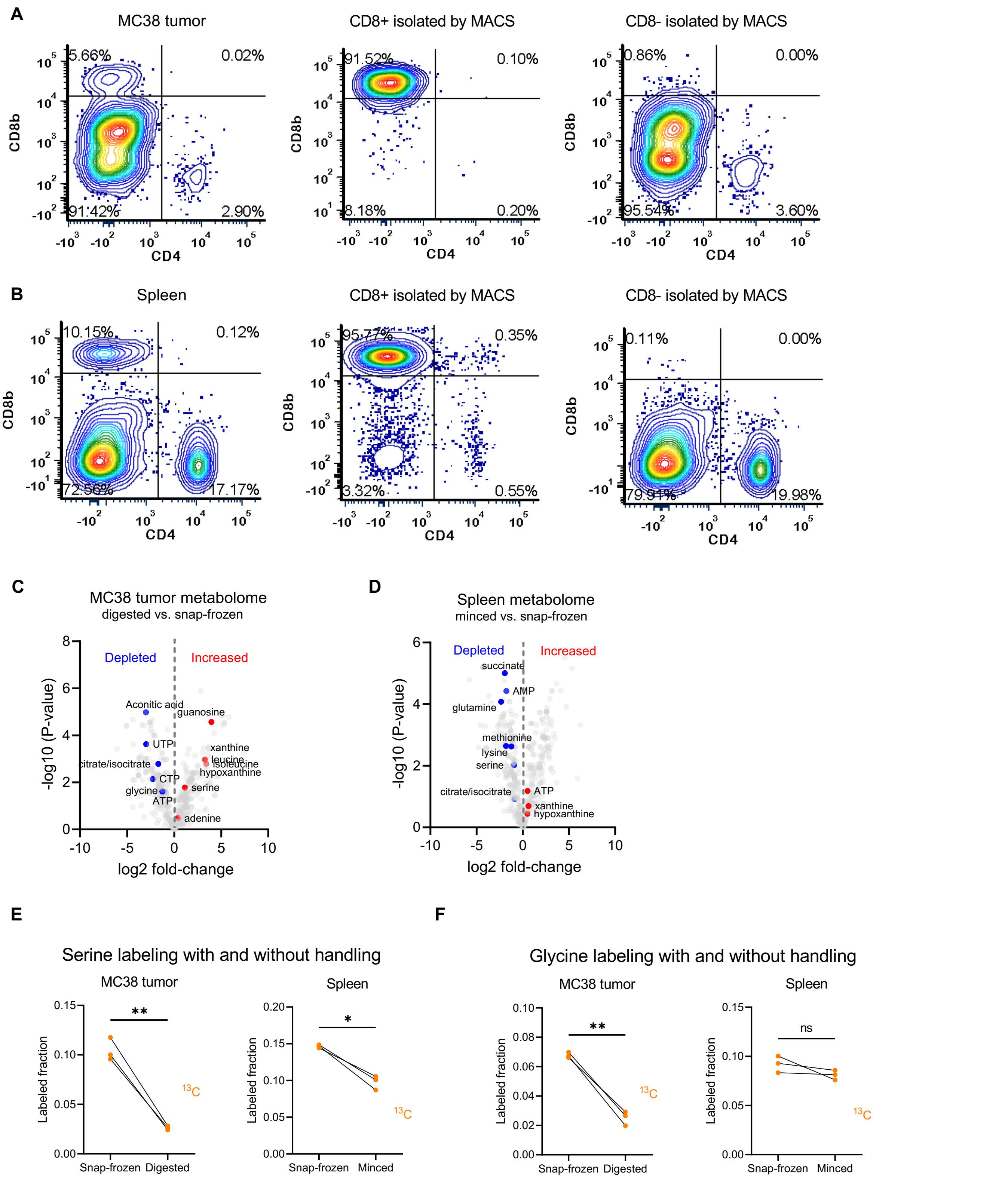


**Supplementary Fig. 2. Cell isolation yielded populations of good purity but altered the abundance and labeling of some metabolites.**

(A–B) Representative flow cytometry result of all cells, $CD8^+$, and $CD8^-$ fractions in (A) MC38 tumor and (B) spleen gated on live $CD45^+$ cells.

(C–F) Changes in metabolite abundance and labeling in dissociated MC38 tumor and spleen compared to the same snap-frozen sample (n = 3).

(C–D) Volcano plots showing changes in metabolite abundance. Metabolite abundance was normalized to the mean of all metabolites (log2 ion count) for each sample.

(E–F) The change of serine (E) and glycine (F) labeling (from [U-$^{13}C$]serine infusion) *P*-values determined by paired two-tailed Student's *t*-tests.

n indicates the number of mice. ns, $p > 0.05$; * $p < 0.05$; ** $p < 0.01$.

Figure S3

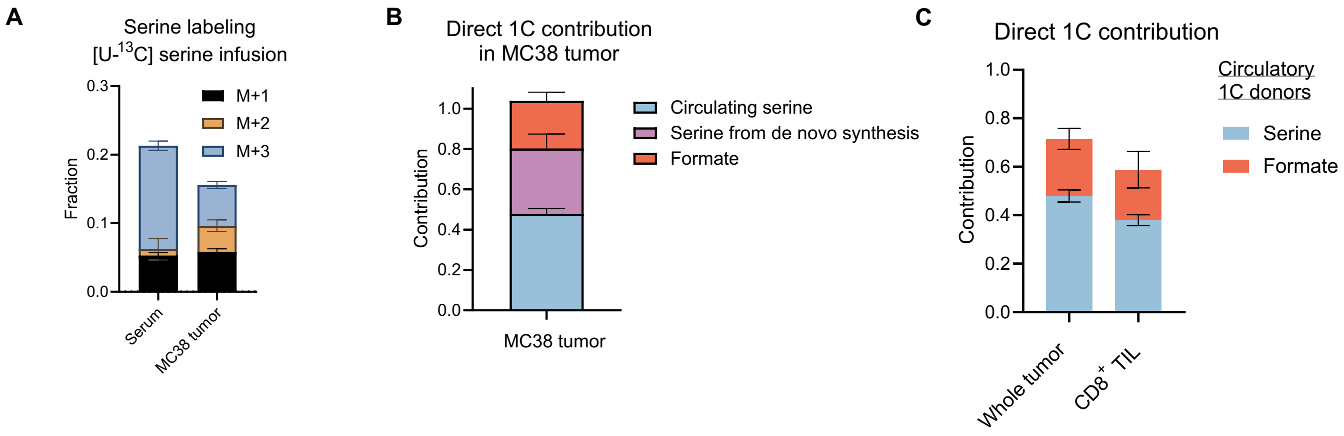


**Supplementary Fig. 3. 1C unit input in MC38 tumor and CD8$^+$ TILs.**
(A) Serine labeling pattern in serum and MC38 tumor for [U-$^{13}$C]serine infusion. Mean ± SD. n = 3 mice.
(B) Direct contribution to tumor 1C units from circulating formate, circulating serine, and serine de novo synthesized in tumor. Mean ± SEM.
(C) Direct contribution of circulating serine and formate to 1C units in bulk MC38 tumor and CD8$^+$ TILs. Mean ± SEM.

**Figure S4**

A

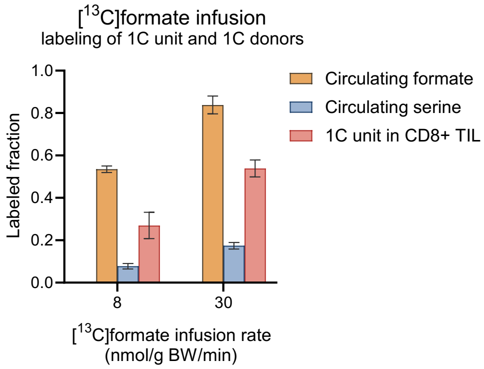


**Supplementary Fig. 4. Elevated circulating formate feeds $CD8^+$ TILs directly.**
(A) Serum serine, serum formate, and 1C unit in $CD8^+$ TILs labeling at different [$^{13}$C]formate infusion rates. TIL labeling markedly exceeds circulating serine labeling, implying that the contribution of formate is direct and not primarily via circulating serine. Mean ± SD. At each infusion rate, n = 4 for serine and n = 3 for others.

**Figure S5**

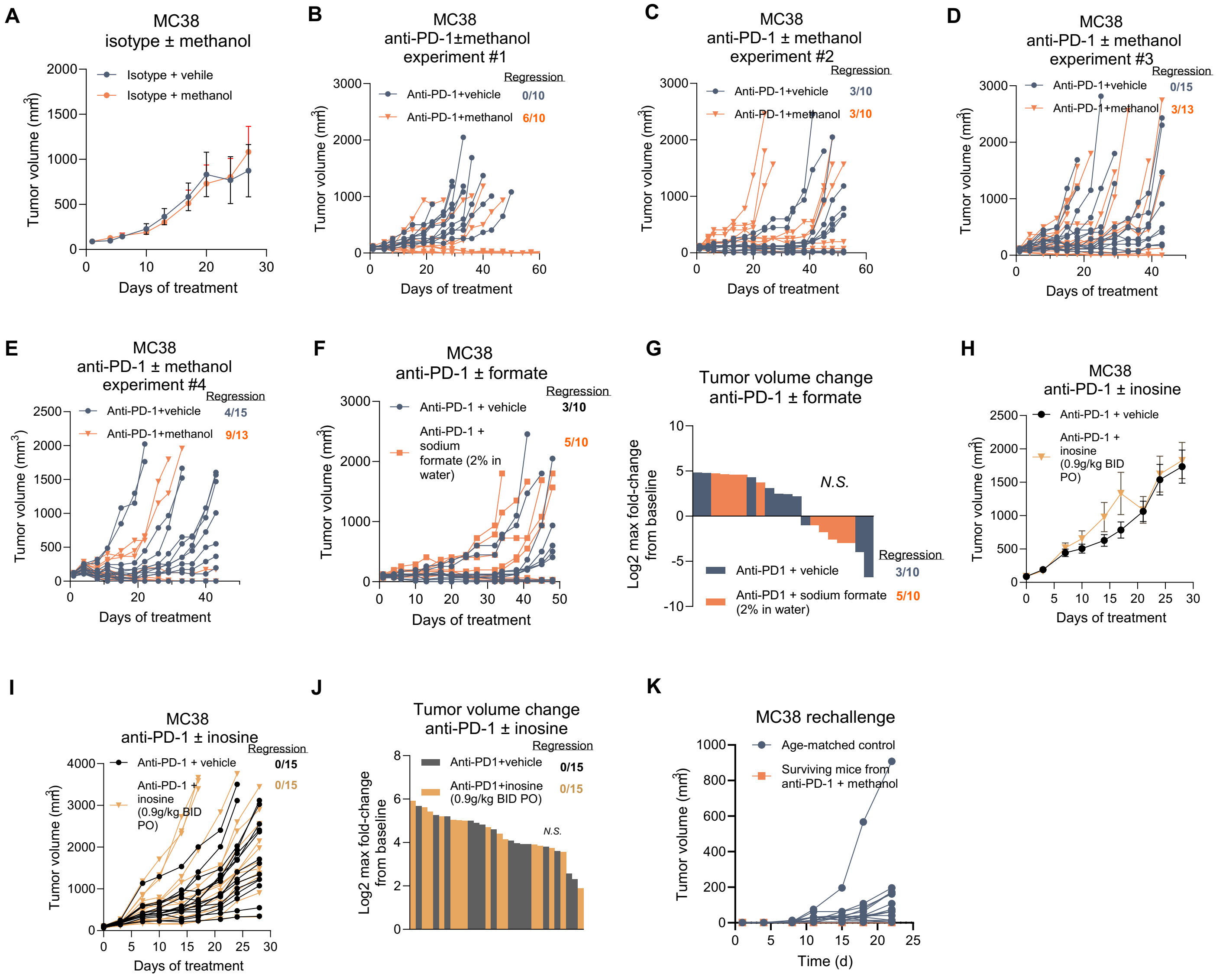


**Supplementary Fig. 5. Effect of supplementing methanol, formate, or inosine on anti-PD-1 efficacy.**
(A) MC38 tumor growth with isotype antibody (5 mg/kg IP, BIW for 2 weeks) ± methanol (2 g/kg daily PO). n = 10.
(B–E) Individual MC38 tumor volumes from 4 independent experiments with anti-PD-1 (5 mg/kg IP, BIW for 2 weeks) ± methanol (2 g/kg daily PO). n is indicated on each panel.
(F–G) MC38 tumor growth with anti-PD-1 (5 mg/kg IP, BIW for 2 weeks) ± sodium formate (2% in drinking water). n = 15.
(F) Individual tumor volumes.
(G) Waterfall plot showing the maximal fold-change of MC38 tumor volume compared to baseline (start of treatment) for each mouse. n = 10 for each group. *P*-value was determined by Pearson chi-square test.
(H–J) MC38 tumor growth with anti-PD-1 (5 mg/kg IP, BIW for 2 weeks) ± inosine (0.9 g/kg BID PO). n = 10.
(H) Average tumor volumes.
(I) Individual tumor volume.
(J) Waterfall plot showing the maximal fold-change of MC38 tumor volume compared to baseline (start of treatment) for each mouse. n = 15. *P*-value was determined by Pearson chi-square test.
(K) Growth of individual tumors after MC38 rechallenge in surviving mice from anti-PD-1 + methanol treatment group (n = 6) and age-matched controls (n = 15).
Mean ± SEM in (A) and (H). n indicates the number of mice.

**Figure S6**

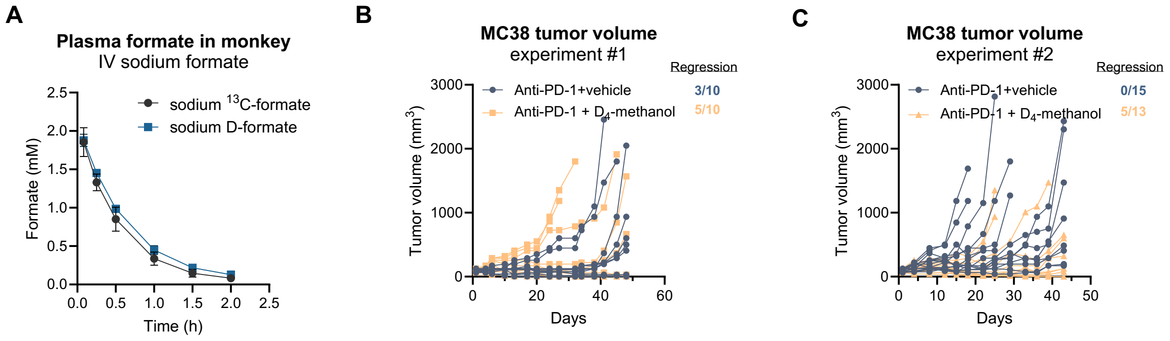


**Supplementary Fig. 6. Pharmacokinetics of deuterated formate and bioactivity of deuterated methanol in combination with checkpoint blockade.**

(A) Plasma formate concentration over time in monkeys intravenously inject with [$^{13}$C] or [D]formate at 50 mg/kg. n = 3.

(B–C) Individual MC38 tumor volumes. Mice were treated with 4 doses of anti-PD-1 with and without daily gavage of 2 g/kg [$D_4$]methanol. n is indicated on each panel.

Mean $\pm$ SD. n indicates the number of animals.
